# A PK-Driven Quantitative Systems Pharmacology Model Predicts Cytokine Release Syndrome Severity Across T Cell-Activating Therapies via a Locked Cytokine Amplification Network

**DOI:** 10.64898/2026.05.05.722920

**Authors:** Hajar Besbassi

## Abstract

Cytokine release syndrome (CRS) is a major dose-limiting toxicity of T cell-engaging immunotherapies. Existing CRS models are drug-class-specific and have not addressed whether a single mechanistic cytokine network can capture severity differences across mechanistically distinct drug classes. Here, we developed a PK-driven quantitative systems pharmacology (QSP) model linking drug exposure, T cell activation dynamics, and a macrophage-amplified cytokine network to clinical CRS severity. The 17-parameter downstream amplification network with macrophage-gated STAT3 positive feedback was developed iteratively. The network was calibrated on blinatumomab, structurally refined using TGN1412 as a transparently disclosed development case, then locked and tested blind on OKT3. The same locked network was used to evaluate cross-drug transferability across three antibody-based T cell engager classes: bispecific, CD28 superagonist, and anti-CD3 with activation-induced cell death. The locked network reproduced the clinically observed CRS severity ordering across all three drugs without re-fitting any shared parameter. The OKT3 blind prediction passed eight qualitative plausibility checks and three of three quantitative cytokine peaks within published clinical ranges. Tocilizumab rescue simulation reproduced five clinically validated phenomena. A mechanistic parameter swap test reversing the T cell exhaustion rate between OKT3 and TGN1412 reversed CRS severity in the expected direction, supporting a mechanistic rather than parameter-fitted interpretation. Local robustness analysis (ABC-style accepted ensemble: 692 of 5,000 parameter sets accepted, 13.8%) and a 2D stability map over the two threshold-setting parameters (0 of 900 wrong-order combinations) confirmed that the cross-drug severity ordering is a property of a feasible parameter region rather than a single tuned point. Profile likelihood analysis of the IL-6 feedback and clearance rates revealed complementary asymmetric profiles consistent with practical identifiability as a ratio. The same locked model predicted three qualitatively distinct dose-response shapes without re-fitting. Findings should be interpreted as a mechanistic proof-of-concept; prospective clinical validation remains pending.

## 1 Introduction

Cytokine release syndrome (CRS) is among the most clinically significant toxicities of T cell-engaging immunotherapies. Bispecific T cell engagers such as blinatumomab cause Grade 1-2 CRS in approximately 15-57% of treated patients [1, 2], while next-generation bispecifics targeting BCMA (teclistamab, talquetamab) and CD20 (mosunetuzumab, glofitamab) extend the class with similar safety challenges. CAR-T cell therapies can trigger severe CRS requiring intensive care management [3], and the management of CAR-T-associated CRS with tocilizumab and corticosteroids has become standard practice across the six FDA-approved CAR-T products. The TGN1412 CD28 superagonist trial in 2006 caused life-threatening CRS in all six healthy volunteers within hours of first dosing [4], an event that fundamentally changed first-in-human trial design and remains the canonical example of catastrophic CRS in modern drug development. OKT3/muromonab-CD3, a murine anti-CD3 antibody historically used in transplant rejection, causes moderate CRS (Grade 2-3) in approximately 95% of patients that is characteristically self-limiting due to activation-induced cell death (AICD) [5, 6]. Despite sharing a common downstream pathway, namely monocyte-derived IL-6 amplification via STAT3 signaling [8], these agents produce CRS of vastly different clinical severity.

CRS now represents a primary dose-limiting toxicity for the rapidly expanding class of T cell-engaging immunotherapies. Severe CRS (Grade 3-4) requires intensive care management, prolongs hospitalization, and drives the dose adjustments and step-dosing protocols used across this drug class. The disparity between the relatively predictable clinical CRS profile of approved bispecifics and the catastrophic outcome of the TGN1412 first-in-human trial points to a translational gap: existing frameworks cannot reliably distinguish between fundamentally different drug mechanisms before dosing humans. Prospective CRS risk stratification for novel agents, particularly for engineered T cell therapies entering first-in-human studies, has been identified by regulatory and academic stakeholders as a priority area for mechanistic modeling support. Empirical PK/PD models trained on existing clinical data extrapolate poorly to mechanistically novel agents, motivating the development of mechanistic frameworks that encode the biology of T cell activation and downstream cytokine amplification rather than fitting drug-specific dose-response curves.

Existing CRS models are drug-class-specific. Published models for CAR-T CRS [3] and bispecific antibody CRS [2] capture the cytokine cascade for individual agents but do not address whether the same cytokine network architecture can explain severity differences across drug classes. QSP models of the tumor-immune interface [9] focus on efficacy rather than safety. To our knowledge, no published model has demonstrated that a single locked mechanistic cytokine network, without changing any shared downstream parameters, can predict CRS severity for mechanistically distinct drugs.

The central challenge is that CRS severity depends on the interaction between drug-specific T cell activation dynamics and shared downstream cytokine amplification. Blinatumomab activates a subset of T cells via CD3 engagement that requires the presence of CD19+ target cells, producing localized and limited T cell activation. TGN1412 causes polyclonal T cell activation via CD28 superagonism, driving sustained T cell expansion without the natural brake of AICD. OKT3 cross-links CD3 to cause polyclonal activation followed by AICD, which limits T cell expansion and thereby caps the cytokine cascade. These three mechanisms should produce qualitatively different CRS severities through the same downstream IL-6 amplification cascade, but this has not been demonstrated computationally with a single locked downstream network. Mechanistic Quantitative Systems Pharmacology (QSP) models are well-suited to this challenge because they can encode drug-specific upstream pharmacology (receptor engagement, T cell activation kinetics, exhaustion dynamics) and shared downstream physiology (cytokine production, macrophage activation, IL-6 amplification) within a single coupled framework. Threshold-type cytokine amplification, in which positive feedback can produce qualitatively different responses to small differences in upstream input, is naturally represented by ODE-based mechanistic models with macrophage-gated feedback structure. If a shared downstream cytokine network architecture is sufficient to explain CRS severity differences across mechanistically distinct drug classes, with only drug-specific upstream T cell parameters changing, then prospective CRS risk assessment for novel agents could in principle be reduced to characterizing the drug’s preclinical T cell activation phenotype, a parameter set that is routinely measured in early development.

Here, a PK-driven mechanistic QSP model demonstrates that a single locked downstream amplification network captures CRS severity differences across three mechanistically distinct antibody-based T cell-activating therapies, reproduces tocilizumab rescue, and produces three qualitatively distinct dose-response shapes (gradual, plateau, switch-like) without re-fitting any shared parameter. The model was developed iteratively: calibrated on blinatumomab using constrained optimization over the most influential subset of cytokine parameters, structurally refined using TGN1412 as a transparently disclosed model development case (not a blind prediction), and then locked and tested on two independent blind challenges, namely OKT3 CRS severity prediction and tocilizumab rescue simulation. Local robustness was assessed through an Approximate Bayesian Computation-style accepted parameter ensemble and a 2D stability map of the two threshold-setting parameters; practical identifiability was assessed via profile likelihood analysis; and a mechanistic parameter swap test reversing the T cell exhaustion rate between OKT3 and TGN1412 confirmed that predicted severity differences reflect drug-specific T cell biology rather than parameter fitting.

## 2 Methods

### 2.1 Software implementation

The CRS model was implemented in Python as a modular package with separate engine, validation, and interface layers, with digitized clinical datasets for the three antibody-based T cell engagers analyzed in this work (blinatumomab, TGN1412, OKT3). All ODE systems were solved using the LSODA adaptive-step integrator with relative tolerance 10^*−*8^ and absolute tolerance 10^*−*10^. Automated tests covered numerical integration, parameter loading, and reproducibility verification.

### 2.2 CRS model: mechanistic cytokine network

The CRS model comprised three coupled subsystems: drug pharmacokinetics, T cell state dynamics, and a five-cytokine network. The combined ODE system contained 11 state variables (Figure 1).

**Figure 1:**
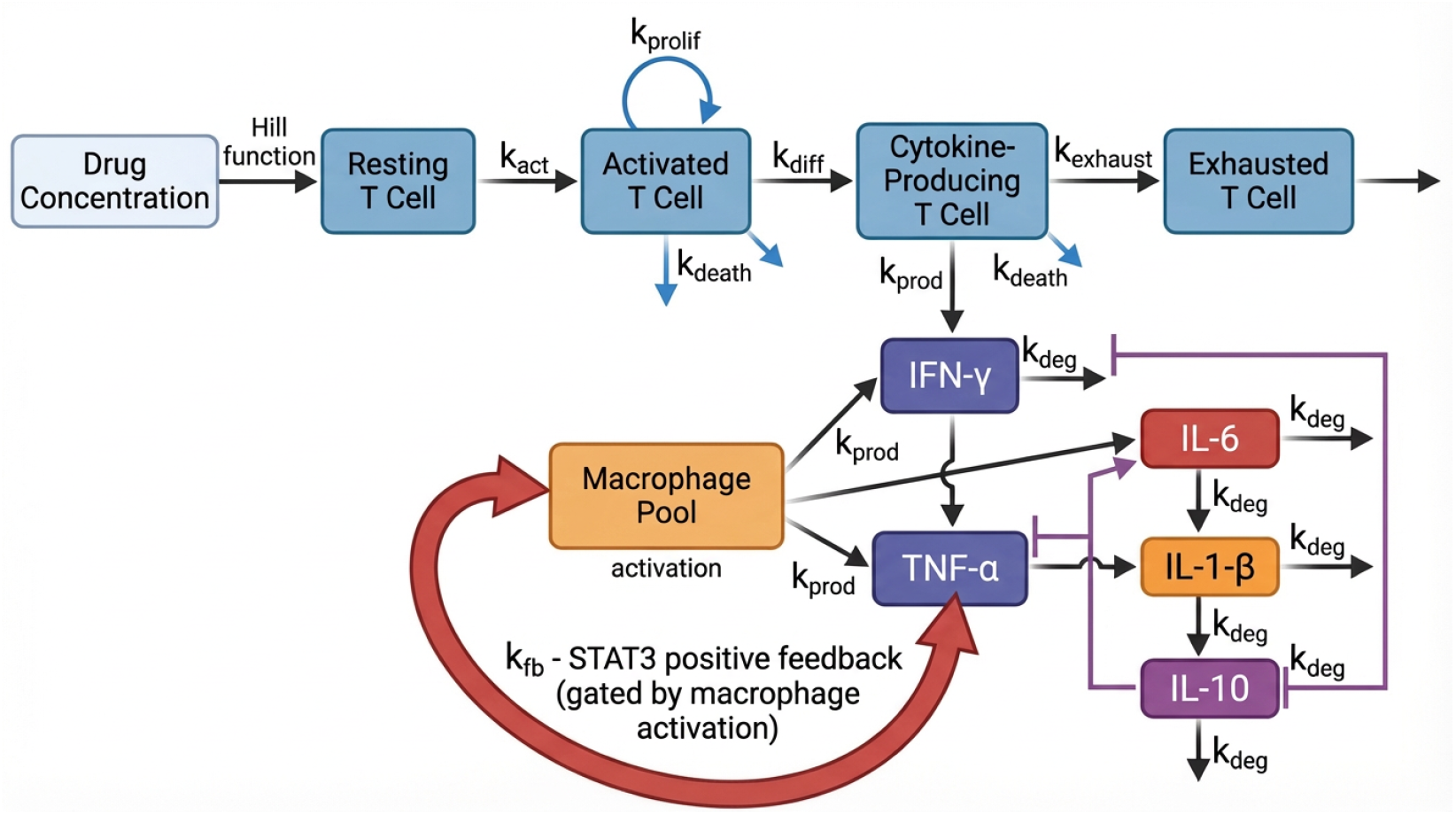
CRS model architecture. Drug concentration activates resting T cells via a Hill function. T cells progress through four states: resting, activated (with proliferation, *k*_*prolif*_), cytokine-producing (*k*_*diff*_), and exhausted (*k*_*exhaust*_). Cytokine-producing T cells release IFN-*γ* and TNF-*α* (*k*_*prod*_), which activate the macrophage pool. Activated macrophages produce IL-6, IL-1*β*, and anti-inflammatory IL-10. IL-6 drives a STAT3 positive feedback loop (*k*_*fb*_, red arrow) gated by macrophage activation. All cytokines degrade at first-order rates (*k*_*deg*_). The 17 shared cytokine network parameters are locked after calibration on blinatumomab; only T cell activation parameters differ between drugs.

#### 2.2.1 Drug pharmacokinetics

A one-compartment model supported both IV bolus and continuous IV infusion:

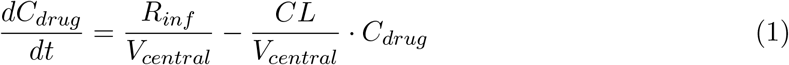

Drug-specific PK parameters are listed in Table 1.

#### 2.2.2 T cell dynamics

Four T cell states were modeled with first-order transitions: resting (*T*_*rest*,0_ = 1,000 cells/*µ*L), activated (*T*_*act*_), cytokine-producing (*T*_*cyt*_), and exhausted (*T*_*exh*_). Drug-driven activation followed a Hill function:

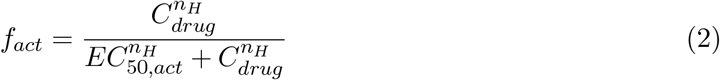

The activation flux was *J*_*act*_ = *k*_*act*_ · *f*_*act*_ · *T*_*rest*_. Activated T cells proliferated with logistic growth (rate *k*_*prolif*_, carrying capacity *T*_*act,max*_), differentiated into cytokine-producing cells (*k*_*diff*_), and exhausted (*k*_*exhaust*_). The exhaustion rate *k*_*exhaust*_ was the key mechanistic differentiator between drug classes: high *k*_*exhaust*_ represented activation-induced cell death (AICD, OKT3), while low *k*_*exhaust*_ represented sustained expansion (TGN1412).

#### 2.2.3 Five-cytokine network with macrophage-gated STAT3 feedback

Five cytokines were modeled: IFN-*γ*, TNF-*α*, IL-6, IL-1*β*, and IL-10. Cytokine-producing T cells produced IFN-*γ* (*k*_*prod,IFNγ*_) and TNF-*α* (*k*_*prod,T NFα*_). These cytokines activated a macrophage pool via a saturating function:

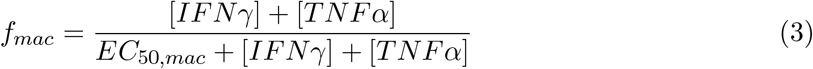

where *EC*_50,*mac*_ = 50 pg/mL. Activated macrophages produced IL-6, IL-1*β*, and the anti-inflammatory cytokine IL-10.

The central mechanistic feature was **macrophage-gated IL-6 STAT3 positive feedback**. Rather than using a constitutive feedback term, the IL-6 amplification rate was gated by the macrophage activation fraction:

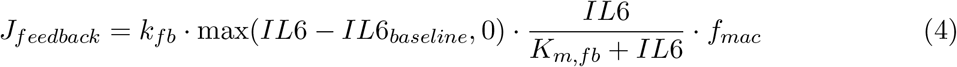

with calibrated values *k*_*fb*_ = 0.528 h^*−*1^ and *K*_*m,fb*_ = 50 pg/mL (Table 2). The macrophage gating (*f*_*mac*_) made the *effective* feedback rate drug-dependent: for blinatumomab (*f*_*mac*_ *≈* 0.3), effective feedback was ∼0.16 h^*−*1^, well below the IL-6 degradation rate (*k*_*deg,IL*6_ = 0.519 h^*−*1^), producing stable IL-6 dynamics. For TGN1412 (*f*_*mac*_ *≈* 0.99), effective feedback approached ∼0.52 h^*−*1^, very close to the degradation rate, entering a near-threshold nonlinear amplification regime that produces substantially higher IL-6 than the other two drug classes. (The term “bifurcation-like” is used descriptively here; a formal local stability or bifurcation analysis of the nonlinear system is left to future work.) This macrophage-gated formulation was motivated by the finding of Norelli and colleagues that monocyte-derived IL-6 is the primary driver of CRS severity [8].

The second key structural feature was **IFN-***γ***-driven macrophage recruitment**. The effective macrophage pool was amplified by inflammatory signals:

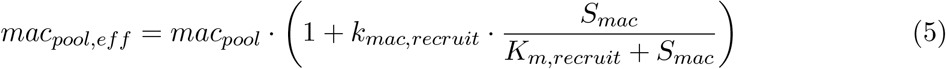

where *S*_*mac*_ = [*IFNγ*] + [*TNFα*], *k*_*mac,recruit*_ = 5.0, and *K*_*m,recruit*_ = 1,000 pg/mL. For blinatumomab (*S*_*mac*_ ∼ 115 pg/mL), *mac*_*pool,eff*_ *≈* 558 (barely changed). For TGN1412 (*S*_*mac*_ > 3,000 pg/mL), *mac*_*pool,eff*_ *≈* 2,500 (5-fold amplification), consistent with the massive monocyte activation observed in the TGN1412 trial [4].

IL-10, produced by activated macrophages, provided anti-inflammatory negative feedback (*IC*_50,*IL*10_ = 50 pg/mL), suppressing all pro-inflammatory cytokine production.

#### 2.2.4 CRS grading

CRS grading in clinical practice follows the ASTCT consensus criteria [12], which are based on clinical symptoms (fever, hypotension, hypoxia) rather than cytokine levels. Because the present model produces cytokine concentrations rather than clinical symptoms, a biomarker-based surrogate grading was used. Peak IL-6 thresholds were selected to approximate the IL-6 ranges associated with each ASTCT clinical grade in published blinatumomab and CAR-T cohorts [2, 11]: Grade 1, IL-6 > 30 pg/mL; Grade 2, IL-6 > 200 pg/mL; Grade 3, IL-6 > 1,000 pg/mL; Grade 4, IL-6 > 5,000 pg/mL. These thresholds were chosen by the author as pragmatic biomarker correlates; they are not clinical grades, are not derived through formal optimization against grade-labeled individual-patient data, and should be interpreted accordingly.

### 2.3 Drug-specific parameterization

The critical design principle was the separation of **locked shared parameters** (the 17-parameter cytokine network, identical across all drugs) from **drug-specific parameters** (PK and T cell activation, which differ by mechanism). Table 1 lists the drug-specific parameters for all three CRS drug classes.

### 2.4 Model development arc: transparent iterative improvement

The CRS model was developed iteratively, and transparency about this process is essential.

#### Step 1: Blinatumomab calibration (formal)

An initial parameter set was chosen by manual adjustment to bring simulated blinatumomab peak IL-6 into the Grade 1-2 range (50-200 pg/mL) reported by Teachey et al. [2]. This initial parameterization was then refined using a formal constrained optimization (Section 2.5) that uses a range-based blinatumomab IL-6 target: the objective returns zero whenever the simulated blinatumomab peak IL-6 lies anywhere within the Teachey 2013 Grade 1-2 range [50, 200] pg/mL, with soft penalties applied to preserve TGN1412 and OKT3 peak IL-6 within their published clinical IL-6 ranges. No specific clinical grade or point-target is forced for any drug. The optimization optimized only the five most influential cytokine network parameters identified by the prior one-at-a-time sensitivity analysis (*k*_*IL*6,*fb*_, *mac*_*pool*_, *k*_*prod,IL*6,*mac*_, *k*_*deg,IL*6_, *EC*_50,*mac*_); the other 12 cytokine network parameters were held fixed. The constrained optimum lies within *±*10% of the initial manual values, indicating that the original manual choices were already in a defensible parameter region.

#### Step 2: TGN1412 failure and structural discovery

The initial model (constitutive IL-6 feedback, *k*_*fb*_ = 0.4 h^*−*1^, no macrophage gating) predicted TGN1412 CRS as Grade 2 (IL-6 = 711 pg/mL), drastically underestimating the clinical Grade 3-4. The root cause was a structural ceiling: with *k*_*fb*_*/k*_*deg,IL*6_ = 0.4*/*0.5 = 0.8 < 1, the IL-6 feedback could never exceed degradation regardless of T cell numbers. Simply increasing *k*_*fb*_ above 0.5 caused *all* drugs to produce Grade 4 CRS, including blinatumomab.

#### Step 3: Macrophage-gated STAT3 feedback

The solution was to make IL-6 feedback conditional on macrophage activation state (Equation 4), consistent with the Norelli 2018 finding [8]. Combined with IFN-*γ*-driven macrophage recruitment (Equation 5), this created a drug-dependent effective feedback rate: low for drugs that weakly activate macrophages (blinatumomab), high for drugs that massively activate macrophages (TGN1412).

**Table 1.** Drug-specific parameters for the three CRS drug classes. The 17 shared cytokine network parameters (Table 2) are identical across all drugs.

| Category | Parameter | Blinatumomab | TGN1412 | OKT3 | Source |
| --- | --- | --- | --- | --- | --- |
| PK | MW (Da) | 54,100 | 148,000 | 150,000 | *a,d,e* |
| | $V_{central}$ (L) | 3.5 | 3.5 | 3.5 | *a,d,e* |
|  | CL (L/h) | 2.1 | 0.005 | 0.02 | *a,d,e* |
| T cell | $EC_{50,act}$ (nM) | 1.0 | 1.0 | 2.0 | *b,f* |
| | $n_H$ (Hill) | 1.5 | 3.0 | 1.5 | *c* |
| | $k_{act}$ ($h^{-1}$) | 0.08 | 0.50 | 0.30 | *f,g* |
| | $k_{prolif}$ ($h^{-1}$) | 0.03 | 0.10 | 0.03 | *g,h* |
| | $k_{exhaust}$ ($h^{-1}$) | 0.005 | 0.012 | 0.08 | *c,i* |
| | $k_{diff}$ ($h^{-1}$) | 0.03 | 0.08 | 0.06 | *f,g* |
| | $T_{act,max}$ (cells/μL) | 5,000 | 20,000 | 5,000 | *j* |
| | Clinical dose | 28 μg/d | 0.1 mg/kg | 5 mg | Approved/trial |
|  | Route | cIV 28d | IV bolus | IV bolus |  |

<sup>a</sup> **Blinatumomab PK.** Clements 2020 [16] popPK: CL = 2.22 L/h, $V_c$ = 5.98 L. Present CL = 2.1 L/h is within reported range; present $V_{central}$ = 3.5 L reflects a one-compartment approximation. Within the 0-72 h CRS window, severity is dominated by T cell activation kinetics, so this does not materially affect predictions.
<sup>b</sup> **In vivo $EC_{50}$ for T cell activation** in the nM range (Klinger 2012 [19]).
<sup>c</sup> **TGN1412 Hill exponent.** $n_H$ = 3.0 reflects cooperative CD28 superagonist signaling (Stebbins 2007 [10]).
<sup>d</sup> **TGN1412 PK.** No human PK published; CL = 0.005 L/h is consistent with typical IgG4 mAb clearance 0.003-0.013 L/h (Dirks 2010 [17]).
<sup>e</sup> **OKT3 PK.** $t_{1/2} \approx 18$ h (Hooks 1991 [18]) implies CL $\approx 0.13$ L/h at typical $V_d$ . Present CL = 0.02 L/h is a known simplification. OKT3 CRS in the model is driven by AICD kinetics (faster timescale than antibody elimination), so predicted grade is expected to be robust to CL values in the 0.02-0.2 L/h range. Formal verification listed in Limitations.
<sup>f</sup> **T cell activation and differentiation rates** taken from QSP T cell engager models (Ma 2020 [22]).
<sup>g</sup> **Proliferation timescale.** Subsequent division time $\approx 9$ h corresponds to $k_{prolif} \approx 0.077$ $h^{-1}$ ; first division 30-50 h (Hawkins 2007 [21]). Present range 0.03-0.10 $h^{-1}$ spans first-to-subsequent division kinetics.
<sup>h</sup> **TGN1412 proliferation.** $k_{prolif} = 0.10$ $h^{-1}$ (doubling time $\sim 7$ h) reflects superagonist-driven sustained expansion without AICD.
<sup>i</sup> **Exhaustion rates.** OKT3 $k_{exhaust} = 0.08$ $h^{-1}$ represents AICD (Chatenoud 1989 [5]); TGN1412 $k_{exhaust} = 0.012$ $h^{-1}$ reflects sustained expansion.
<sup>j</sup> **T cell carrying capacity.** $T_{act,max}$ is a model-assumed upper bound constraining activated T cell expansion; normal lymphocyte range is 1,000-4,800 cells/ $\mu$ L. Values of 5,000-20,000 cells/ $\mu$ L were assumed to reflect polyclonal activation capacity and were not directly measured.

**Table 2.** Locked cytokine network parameters. These 17 parameters are identical across all three drug classes and across the tocilizumab rescue simulation.

| Parameter | Symbol | Value | Unit |
| --- | --- | --- | --- |
| IFN-γ production rate | $k_{prod,IFN\gamma}$ | 0.30 | pg/mL/h per cell/μL |
| TNF-α production rate | $k_{prod,TNF\alpha}$ | 0.15 | pg/mL/h per cell/μL |
| IL-6 macrophage production rate* | $k_{prod,IL6,mac}$ | 0.108 | pg/mL/h per cell/μL |
| Baseline macrophage pool* | $mac_{pool}$ | 465 | cells/μL equiv. |
| IL-6 STAT3 feedback gain* | $k_{fb}$ | 0.528 | $h^{-1}$ |
| IL-6 feedback $K_m$ | $K_{m,fb}$ | 50 | pg/mL |
| Macrophage activation $EC_{50}$* | $EC_{50,mac}$ | 48.6 | pg/mL |
| IL-6 degradation rate* | $k_{deg,IL6}$ | 0.519 | $h^{-1}$ |
| TNF-α degradation rate | $k_{deg,TNF\alpha}$ | 0.70 | $h^{-1}$ |
| IFN-γ degradation rate | $k_{deg,IFN\gamma}$ | 0.30 | $h^{-1}$ |
| IL-1β degradation rate | $k_{deg,IL1\beta}$ | 0.40 | $h^{-1}$ |
| IL-10 degradation rate | $k_{deg,IL10}$ | 0.35 | $h^{-1}$ |
| IL-10 suppression $IC_{50}$ | $IC_{50,IL10}$ | 50 | pg/mL |
| IL-1β macrophage production | $k_{prod,IL1\beta,mac}$ | 0.002 | pg/mL/h per cell/μL |
| IL-10 macrophage production | $k_{prod,IL10,mac}$ | 0.01 | pg/mL/h per cell/μL |
| Macrophage recruitment gain | $k_{mac,recruit}$ | 5.0 | - |
| Recruitment half-saturation | $K_{m,recruit}$ | 1,000 | pg/mL |

\* **Formally calibrated.** Parameter values were determined by formal constrained optimization against published blinatumomab clinical IL-6 data (Section 2.5). The other 12 cytokine network parameters were held fixed during optimization.
**Degradation rates.** Reflect effective cytokine decay in the lumped compartmental model, corresponding to in-model half-lives of $\sim 1$ -2 h for pro-inflammatory cytokines. Within the order-of-magnitude range of plasma half-lives reported for TNF-α ( $\sim 0.5$ h at therapeutic doses; Ferraiolo 1989 [20]) and IL-6 (1-4 h in the absence of soluble receptor binding; Nishimoto 2008 [14]), and comparable to effective degradation rates used in previous QSP cytokine models (Weddell 2023 [15]).
**Macrophage pool.** Baseline $mac_{pool}$ reflects peripheral monocyte counts in the normal range (200-800 cells/ $\mu$ L) plus tissue-resident macrophages in a lumped representation.
**Identifiability.** Formal parameter identifiability analysis for the 17-parameter network is listed as a priority in Limitations. Profile likelihood for the two threshold-setting parameters is reported in Section 3.10.

#### Step 4: Parameter lock and blind testing

Following re-calibration, the 17 cytokine network parameters were locked and tested on OKT3 and tocilizumab rescue without any changes.

##### TGN1412 is not a blind prediction

The model structure was improved because of TGN1412 failure. This is standard iterative model development in QSP (calibrate, test, fail, improve, retest), but it must be distinguished from the OKT3 and tocilizumab results, which are true blind predictions from the locked model.

##### Exploratory CAR-T transfer test (Supplementary Section S2)

The locked downstream cytokine network was also applied, as a post hoc exploratory transfer test, to tisagenlecleucel (CTL019/Kymriah) CAR-T cell therapy by replacing the drug pharmacokinetic block with a CAR-T cell expansion module. This is not a blind validation: CAR-T-specific upstream parameters were formulated with knowledge of the Teachey 2016 [3] cytokine ranges. The 17 cytokine network parameters remained locked at the blinatumomab-calibrated values. CAR-T is not part of the cross-drug calibration or the main antibody-based claim. Full methods, equations, parameter choices, a transfer-test table, and a figure are provided in Supplementary Section S2.

### 2.5Formal calibration procedure

To strengthen the credibility of the locked cytokine network parameters, a formal constrained optimization was performed against published blinatumomab clinical data while preserving the cross-drug separability that defines the architectural transferability claim of this work.

#### Calibration target

The primary target is range-based: blinatumomab peak IL-6 must fall within the Grade 1-2 IL-6 range 50-200 pg/mL reported by Teachey et al. [2] for blinatumomab CRS. The objective returns zero whenever the simulated blinatumomab peak IL-6 lies anywhere within this range; no specific point value within the range is forced. The same source was cited in the original manual calibration.

#### Free parameters

Five cytokine network parameters were treated as free, selected based on the prior one-at-a-time sensitivity analysis as the top contributors to peak IL-6 (Section 4, Figure 10): *k*_*IL*6,*fb*_, *mac*_*pool*_, *k*_*prod,IL*6,*mac*_, *k*_*deg,IL*6_, and *EC*_50,*mac*_. The other 12 cytokine network parameters were held fixed at their initial values, and all drug-specific T cell parameters were also held fixed (only the shared cytokine network was calibrated).

#### Objective function

The objective combines a range-based blinatumomab IL-6 term with soft cross-drug architectural preservation penalties. Define a range-based log-fold error

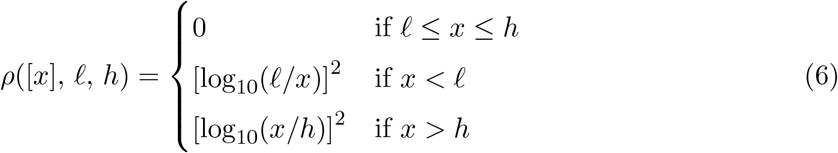

that returns zero whenever *x* lies within [*f, h*] and otherwise returns the squared log-fold distance to the nearest boundary. Then

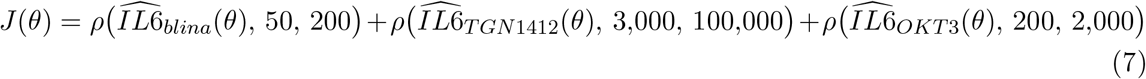

with all three IL-6 ranges taken a priori from the published clinical sources cited in this paper (Teachey 2013 for blinatumomab, Suntharalingam 2006 for TGN1412, Chatenoud 1989 / Abramowicz 1989 / Ferran 1990 for OKT3). A hard rejection (*J* = 10^6^) is applied if any drug undergoes IL-6 amplification runaway (peak IL-6 > 10^6^ pg/mL).

#### No specific clinical grade is required for any drug; the objective only requires each prediction to fall inside the published clinical IL-6 range

In particular, TGN1412 is not forced to the Grade 4 threshold; clinical TGN1412 is described as “Grade 3-4 in all six volunteers” [4], so any prediction within the 3,000-100,000 pg/mL range is acceptable. The objective is therefore best described as a **constrained blinatumomab calibration with soft cross-drug architectural preservation constraints**, not as a pure single-drug fit. The calibration uses blinatumomab IL-6 as its primary range target; TGN1412 and OKT3 information enters only through the architectural preservation penalties (which require their predictions to remain in published clinical IL-6 ranges) and not through any cytokine-magnitude fit.

#### Optimizer

Derivative-free Nelder-Mead optimization starting from the initial manual values. Bounds were *±*50% of the initial values, applied to keep the optimization in a smooth region away from the STAT3 amplification boundary. Convergence was verified by inspection of the objective trajectory and the function evaluation count.

#### Result

The optimization converged with *J* (*θ*_*opt*_) = 0 (machine precision), meaning that the calibrated parameter set produces blinatumomab peak IL-6 inside [50, 200] pg/mL, TGN1412 peak IL-6 inside [3,000, 100,000] pg/mL, and OKT3 peak IL-6 inside [200, 2,000] pg/mL simultaneously, with no residual log-fold error against any of the three range boundaries. Starting the optimizer from the original manual values, all parameter changes were less than *±*10%, and re-running the optimizer from the calibrated point produced trivial movements (< 1.1% change in any parameter), confirming that the calibrated point sits at a local minimum of the objective. The final calibrated parameters used in this work are documented in Table 2.

#### Robustness check

A local stability check perturbed each optimized parameter by *±*10% individually and recomputed the objective. Three of the five parameters (*mac*_*pool*_, *k*_*prod,IL*6,*mac*_, *EC*_50,*mac*_) showed objective changes below 0.02, indicating local insensitivity. Two parameters (*k*_*IL*6,*fb*_ and *k*_*deg,IL*6_) showed asymmetric responses: small perturbations in the runaway-amplification direction triggered the hard rejection criterion, confirming that the calibrated point sits near the boundary of the IL-6 STAT3 amplification regime. This is biologically meaningful and is consistent with the sensitivity analysis observation (Section 4) that the model is intentionally placed near the boundary between self-limiting and catastrophic CRS.

#### Scope and interpretation

This is a deliberately modest constrained optimization with a range-based target on a single cytokine peak, a small free parameter set (5 of 17), bounded parameter ranges, and soft cross-drug architectural preservation constraints. It is not a multiobjective fit of all cytokines. Its purpose is to convert the original manual calibration into a procedure with an explicit objective function, an explicit free parameter set, explicit constraints, and an explicit convergence criterion, all of which can be reproduced from the published code and parameter targets.

#### Local accepted parameter ensemble around the calibrated region

To assess whether the calibrated parameter point is an isolated optimum or sits within a robust feasible region, an ABC-style local accepted parameter ensemble was constructed around the calibrated values. This is a **local** robustness analysis, not a full Bayesian posterior characterization: the prior is uniform within *±*30% of the calibrated point on each of the five free parameters, the acceptance rule is range-based on a single output (peak IL-6 only), and no proper likelihood function or MCMC convergence is computed. The purpose is to characterize the size and shape of the feasible region in a finite neighborhood of the calibrated point, not to explore the full plausible biology.

##### Acceptance ranges (defined a priori from published clinical literature, held fixed throughout)

A candidate parameter set is accepted if and only if its predicted peak IL-6 falls simultaneously within all three published clinical ranges:

- blinatumomab 50-200 pg/mL (Teachey 2013, Grade 1-2) [2]
- TGN1412 3,000-100,000 pg/mL (Suntharalingam 2006) [4]
- OKT3 200-2,000 pg/mL (Chatenoud 1989, Abramowicz 1989, Ferran 1990) [5, 6, 7]

These ranges were taken from the published clinical sources cited in the calibration design and were not modified after observing the ensemble results. Candidates that triggered IL-6 amplification runaway (peak > 10^6^ pg/mL) were also rejected as non-physical. No specific clinical grade was required for any drug; only that the predicted peak IL-6 fall within the published range.

##### Important scope notes

1. **Acceptance is IL-6-only**. The acceptance rule checks peak IL-6 only. The other cytokines in the network (IFN-*γ*, TNF-*α*, IL-10, IL-1*β*) are simulated and tracked diagnostically but are not included in the acceptance criterion. The ensemble therefore characterizes the robustness of the **IL-6-centered severity architecture**, not the robustness of the full multi-cytokine network. Robustness of the non-IL-6 cytokines remains a target for future calibration work.
2. **The procedure involves only the three antibody-based drugs**. The ABC ensemble, the formal calibration, and the stability map all use blinatumomab, TGN1412, and OKT3 only.
3. **The prior is local, not biological**. The *±*30% bounds reflect a robustness neighborhood around the calibrated values, not a wider biologically motivated parameter space. Conclusions about feasibility apply within this neighborhood; broader biological identifiability would require an informed prior and a proper Bayesian or profile-likelihood treatment.

#### Multi-cytokine acceptance test

To test whether the locked cytokine network is robust on more than the IL-6 axis, the same 5,000-sample local prior was re-evaluated against an extended acceptance rule that adds five further cytokine constraints from published clinical sources (TGN1412 IFN-*γ* in 1,000-5,000 pg/mL and TNF-*α* in 1,000-3,500 pg/mL from Suntharalingam 2006 [4]; OKT3 IFN-*γ* in 50-500 pg/mL from Ferran 1990 [7] and TNF-*α* in 100-800 pg/mL from Abramowicz 1989 [6] / Ferran 1990 [7]). Blinatumomab cytokines other than IL-6 are not consistently reported in Teachey 2013 case-series and were left out of the rule to keep it fair. The full rule therefore checks 7 cytokine-by-drug constraints simultaneously, all defined a priori from the published clinical literature. A second variant that drops the single failing constraint (TGN1412 TNF-*α*) is also reported transparently as a conditional sensitivity analysis.

#### Practical identifiability via profile likelihood

To assess whether the two threshold-setting parameters *k*_*IL*6,*fb*_ and *k*_*deg,IL*6_ were practically identifiable as individual parameters, profile-likelihood analyses were performed for each. For each target parameter, a 21-point grid spanning *±*30% of the calibrated value was evaluated. At each fixed grid value, the remaining four free parameters were re-optimized using the same constrained calibration objective, Nelder-Mead optimizer, and *±*50% bounds defined above, yielding the minimum achievable objective value *J* (*θ*) at each grid point. The resulting objective-parameter profiles distinguish practical identifiability, indicated by a sharp U-shaped profile, from compensatory non-identifiability, indicated by a relatively flat profile in which other parameters absorb the imposed change. This analysis should be interpreted as a local profile-based assessment of practical identifiability, not a full global identifiability analysis, which would require a broader biologically informed prior, parameter-correlation analysis, and ideally a Bayesian or rank-based identifiability framework.

#### Cross-drug stability map

To characterize the local parameter-region structure, a 2D stability map was generated over the two parameters most directly involved in the IL-6 STAT3 amplification balance: *k*_*IL*6,*fb*_, the feedback rate, and *k*_*deg,IL*6_, the IL-6 clearance rate. A 30 *×* 30 grid spanning *k*_*IL*6,*fb*_ *∈* [0.35, 0.75] h^*−*1^ and *k*_*deg,IL*6_ *∈* [0.35, 0.75] h^*−*1^ was evaluated. At each grid point, the three antibody-based drugs blinatumomab, TGN1412, and OKT3 were simulated. Each grid point was classified as: (i) **Runaway** if any drug crossed the IL-6 amplification boundary (> 10^6^ pg/mL); (ii) **Wrong order** if the cross-drug severity ordering was disrupted, defined as blinatumomab *≥* OKT3 or OKT3 *≥* TGN1412; (iii) **Out of range** if at least one drug fell outside its clinical IL-6 range; or (iv) **All in range** if all three drugs fell within their clinical IL-6 ranges. The same a priori clinical ranges as the local accepted ensemble were used. The other three free parameters were held fixed at their calibrated values during the sweep.

### 2.6 Tocilizumab rescue simulation

To test whether the locked model could predict therapeutic intervention, tocilizumab (anti-IL-6R) rescue of TGN1412-severity CRS was simulated. The simulation used a two-phase ODE approach:

**Phase 1** (0 to *t*_*intervention*_): Normal CRS dynamics as described above.

**Phase 2** (*t*_*intervention*_ to *t*_*end*_): IL-6R blockade was modeled by two modifications to the ODE system:

- STAT3 feedback set to zero (*J*_*feedback*_ = 0), reflecting complete IL-6R blockade
- IL-6 degradation reduced to 30% of normal (*k*_*deg,IL*6,*eff*_ = 0.3 · *k*_*deg,IL*6_), reflecting loss of receptor-mediated IL-6 clearance

All other equations (T cell dynamics, IFN-*γ* production, TNF-*α* production, macrophage activation) remained unchanged, reflecting the IL-6-independent nature of these pathways. Thirteen intervention timepoints were tested: 0, 1, 2, 3, 4, 6, 8, 12, 18, 24, 36, 48, and 72 h.

No parameters were changed from the locked model. The tocilizumab pharmacology was implemented as a structural modification (receptor blockade), not a parameter change.

### 2.7 Dose-response analysis

Dose-response sweeps were performed for all three drugs using the locked model:

1. Blinatumomab: 10 doses (0.001-0.1 mg, single-dose bolus equivalents)
2. TGN1412: 10 doses (0.0001-0.5 mg/kg)
3. OKT3: 9 doses (0.5-20 mg)

At each dose, a single-patient simulation was run (70 kg body weight), and CRS grade and peak IL-6 were recorded. No parameters were changed between doses.

### 2.8 Mechanistic parameter swap test

To distinguish between real mechanistic predictions and arbitrary parameter fitting, the exhaustion rate constant (*k*_*exhaust*_) was swapped between OKT3 and TGN1412:

1. OKT3 with TGN1412’s *k*_*exhaust*_ (0.012 h^*−*1^): removes AICD
2. TGN1412 with OKT3’s *k*_*exhaust*_ (0.08 h^*−*1^): forces AICD

If the model captures a real biological distinction (AICD vs. sustained expansion), swapping this single parameter should reverse CRS severity.

### 2.9 Virtual population generation

Virtual populations (*N* = 100 per drug) were generated by sampling inter-individual variability (IIV) on selected model parameters. Five T cell activation parameters (*k*_*act*_, *k*_*prolif*_, *k*_*exhaust*_, *EC*_50,*act*_, *T*_*act,max*_), the baseline macrophage pool size (*mac*_*pool*_), and body weight were sampled independently from log-normal distributions with coefficient of variation (CV) = 25-30%. These CV values were selected based on typical IIV ranges reported for immune cell parameters in QSP models [22] and pharmacokinetic variability in biologics [17]. Physiological covariates (weight, height, BSA, eGFR) were sampled with allometric correlations; drug clearance and volume of distribution were scaled to body weight.

No acceptance criteria were applied to filter virtual patients; all sampled parameter sets were retained. This approach provides an illustrative assessment of population-level CRS grade distributions under assumed IIV, rather than a formally calibrated virtual population matched to patient-level data. The absence of individual patient-level cytokine time-course data for calibration of IIV precludes rigorous virtual population validation at this stage. Population results should therefore be interpreted as sensitivity to parameter variability rather than as quantitative predictions of clinical grade distributions.

### 2.10 Computational environment

Simulations were performed in Python 3.11 with standard scientific computing libraries. A reproducibility test suite of 67 automated tests covered numerical integration, parameter loading, and end-to-end simulation outputs.

## 3 Results

### 3.1 Three-drug CRS validation

Figure 2 provides a graphical overview of the mechanistic architecture and the three CRS outcomes produced by the locked cytokine network.

The locked cytokine network reproduced clinically observed CRS severity differences across the three mechanistically distinct antibody-based T cell engagers (Table 3, Figure 3). All simulations used identical shared cytokine network parameters; only drug-specific PK and T cell activation parameters differed. Predicted CRS grades increased from blinatumomab (Grade 1, peak IL-6 ∼100 pg/mL) through OKT3 (Grade 2, peak IL-6 671 pg/mL) to TGN1412 (Grade 3, peak IL-6 4,349 pg/mL), corresponding to a ∼43-fold span in peak IL-6 across the three drugs, with each prediction falling within the published clinical IL-6 range for the corresponding agent. The severity ordering therefore emerged from differences in upstream PK and T cell activation alone, propagating through one shared downstream amplification network.

**Figure 2:**
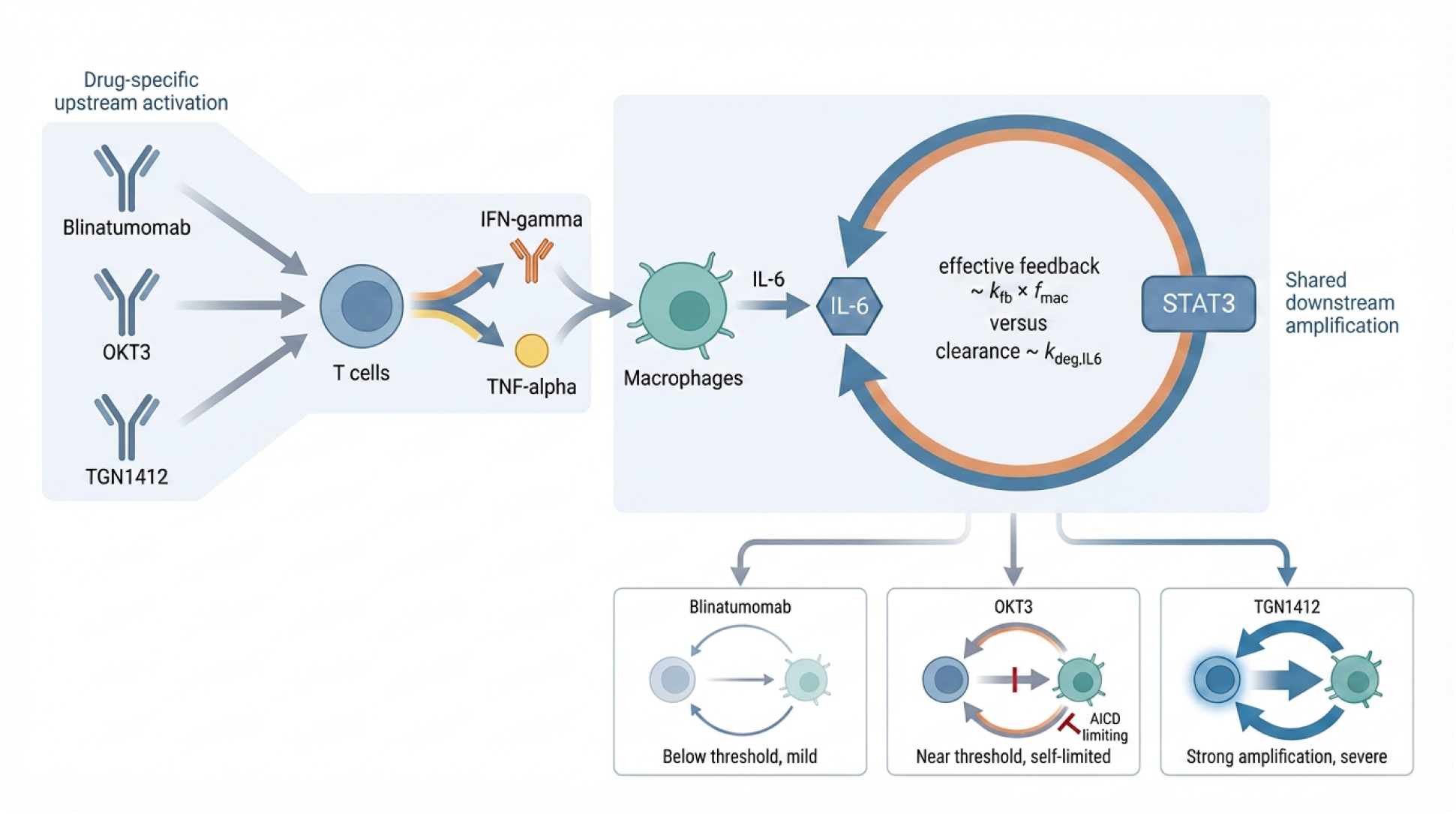
Conceptual overview of the cross-drug CRS mechanism. Left: drug-specific upstream T cell activation. Each of the three antibody-based T cell engagers activates T cells with characteristic kinetics. Blinatumomab (CD19 *×* CD3 bispecific) activates a limited subset of T cells in the presence of CD19+ targets. OKT3 (anti-CD3) activates many T cells, but the response is self-limited by activation-induced cell death (AICD). TGN1412 (CD28 superagonist) drives broad T cell activation without the AICD brake. Right: shared downstream amplification network. Activated T cells produce IFN-*γ* and TNF-*α*, which activate macrophages. Macrophages produce IL-6, and IL-6 reinforces its own production through macrophage-gated STAT3 positive feedback. The effective feedback rate *k*_*IL*6,*fb*_ · *f*_*mac*_ versus the IL-6 clearance rate *k*_*deg,IL*6_ determines whether the system remains controlled or enters strong amplification. Bottom: the same locked downstream network produces three qualitatively distinct operating points: blinatumomab below threshold (mild), OKT3 closer but capped by AICD (moderate), and TGN1412 in the strong-amplification regime (severe).

**Table 3.** Three-drug CRS comparison from one locked cytokine network.

| Drug | Mechanism | CRS Grade<br>(predicted) | Peak cytokines (pg/mL) |  |  | Status |
| --- | --- | --- | --- | --- | --- | --- |
| | | | IL-6 | TNF-α | IFN-γ | |
| Blinatumomab | BiTE (CD19×CD3) | 1 | ~100 | ~63 | ~9 | Calibration |
| TGN1412 | CD28 superagonist | 3 | 4,349 | 558 | 1,674 | Development |
| OKT3 | Anti-CD3 (AICD) | 2 | 671 | 174 | 207 | **Blind** |

**Blinatumomab.** Calibration drug. Predicted Grade 1, consistent with clinical Grade 1-2 reported in Teachey 2013 [2].
**TGN1412.** Model development case, not a blind prediction. Predicted Grade 3 with peak IL-6 = 4,349 pg/mL, within clinical Grade 3-4 and the 3,000-100,000 pg/mL range reported by Suntharalingam 2006 [4].
**OKT3.** True blind prediction. Predicted Grade 2 with peak IL-6 = 671 pg/mL, within clinical Grade 2-3 reported by Chatenoud 1989 [5].

**Figure 3:**
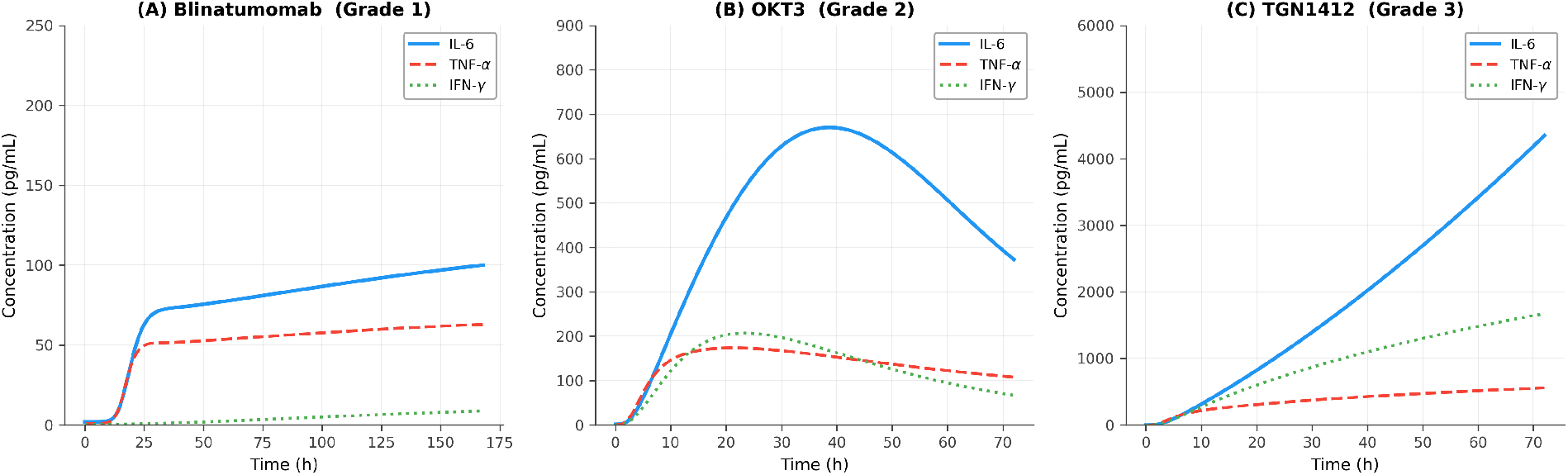
CRS cytokine dynamics across three drug classes from one locked model. IL-6 (solid), TNF-*α* (dashed), and IFN-*γ* (dotted) time-courses for blinatumomab (A, Grade 1), OKT3 (B, Grade 2), and TGN1412 (C, Grade 3). Note different y-axis scales reflecting the ∼43-fold IL-6 difference between blinatumomab (100 pg/mL) and TGN1412 (4,349 pg/mL). All three simulations used identical cytokine network parameters; only drug-specific PK and T cell activation parameters differed (Table 1).

### 3.2 TGN1412 model development case

TGN1412 was used as a model development case rather than a blind validation. The macrophage-gated STAT3 feedback structure was introduced specifically to address the initial model’s failure to reproduce the TGN1412 cytokine profile (Section 2.4), and the resulting performance is reported here for completeness.

The model predicted Grade 3 CRS for TGN1412 at 0.1 mg/kg (7 mg in a 70-kg subject), with peak IL-6 = 4,349 pg/mL. Clinical data from the Suntharalingam 2006 trial [4] report Grade 3-4 CRS in all six volunteers, with peak IL-6 ranging from 3,000 to >100,000 pg/mL. The predicted value lies near the lower end of this broad clinical range and is consistent with the observed severe CRS phenotype. The model also reproduced the absence of severe CRS at the Theralizumab dose (0.0001 mg/kg), predicting a mild response (Grade 1, peak IL-6 = 71 pg/mL).

An illustrative population simulation (*N* = 100, see Methods for IIV assumptions) predicted all virtual subjects in Grade 3 (67%) or Grade 4 (33%), consistent with the 6/6 clinical Grade 3-4 incidence (Figure 5). Two known discrepancies should be noted: peak TNF-*α* was underestimated (558 vs. clinical 1,000-3,500 pg/mL), and CRS onset was delayed (Grade 1 at 3 h vs. clinical symptoms at 60-90 min). Table 4 summarizes the comparison.

**Table 4.**
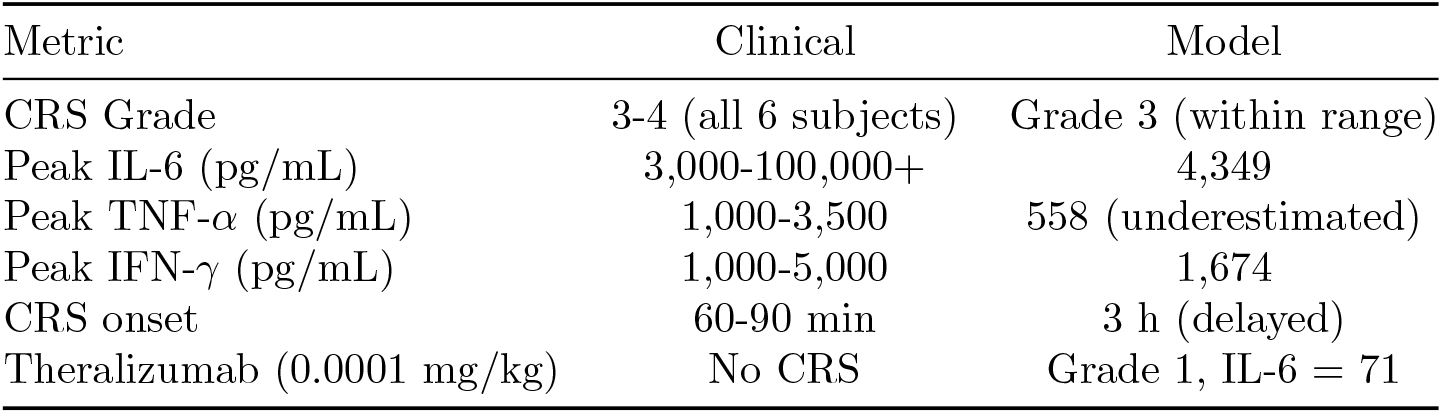
TGN1412 validation against Suntharalingam 2006 clinical data.

### 3.3 OKT3 blind prediction

OKT3/muromonab-CD3 represents a fundamentally different T cell activation mechanism from both blinatumomab and TGN1412. Anti-CD3 cross-linking causes polyclonal T cell activation followed by AICD: T cells activate, then die, limiting the cytokine cascade. This is captured in the model by a high *k*_*exhaust*_ = 0.08 h^*−*1^ and low *T*_*act,max*_ = 5,000 cells/*µ*L, reflecting the self-limiting nature of anti-CD3 activation.

The locked model predicted Grade 2 CRS for OKT3 at 5 mg IV, with peak IL-6 = 671 pg/mL. Two complementary validation approaches were applied. First, eight qualitative clinical plausibility checks (correct CRS grade, cytokine peaks above clinically observed thresholds, onset within 6 h, severity ordering relative to TGN1412) were evaluated against published clinical data (Chatenoud 1989 [5], Abramowicz 1989 [6], Ferran 1990 [7]); all eight passed (Table 5). Second, quantitative cytokine-level comparison against reported clinical ranges is presented in Table 10: all three OKT3 cytokine peaks (IL-6, TNF-*α*, IFN-*γ*) fell within the reported clinical ranges and passed the 2-fold absolute fold-error criterion.

**Table 5.** OKT3 blind prediction: 8/8 validation checks passed.

| Check | Criterion | Clinical | Predicted | Status |
| --- | --- | --- | --- | --- |
| 1 | CRS Grade $\geq 2$ | Grade 2-3 typical | Grade 2 | PASS |
| 2 | CRS Grade $\leq 3$ (not Grade 4) | Not Grade 4 | Grade 2 | PASS |
| 3 | Peak IL-6 $\geq 100 \text{ pg/mL}$ | 200-2,000 | 671 | PASS |
| 4 | Peak IL-6 $\leq 3,000 \text{ pg/mL}$ | <TGN1412 level | 671 | PASS |
| 5 | Peak TNF-$\alpha \geq 50 \text{ pg/mL}$ | 100-800 | 174 | PASS |
| 6 | Peak IFN-$\gamma \geq 50 \text{ pg/mL}$ | 50-500 | 207 | PASS |
| 7 | CRS onset within 6 h | 1-2 h symptoms | 4 h | PASS |
| 8 | IL-6 < TGN1412 IL-6 | Less severe | 671 vs. 4,349 | PASS |

Clinical references: Chatenoud 1989 [5], Abramowicz 1989 [6], Ferran 1990 [7]. Model parameters were locked; zero changes were made to the cytokine network.

The mechanistic basis of the correct prediction was activation-induced cell death (Figure 4). T cell dynamics showed a characteristic peak-and-decline pattern for OKT3: *T*_*cyt*_ peaked at 334 cells/*µ*L at 21 h then declined to 183 cells/*µ*L by 48 h (AICD). In contrast, TGN1412 showed continuously rising *T*_*cyt*_ reaching 2,790 cells/*µ*L at 48 h (sustained expansion). This T cell dynamic difference, propagated through the shared cytokine network, produced a ∼6.5-fold IL-6 difference between the two anti-CD3 mechanisms (OKT3 671 vs. TGN1412 4,349 pg/mL). An illustrative population simulation (*N* = 100, log-normal IIV with CV = 25-30% on T cell activation parameters, macrophage pool size, and body weight; see Methods) was consistent with the single-patient result: ∼96% of virtual patients predicted Grade 2 CRS with ∼4% Grade 3, qualitatively consistent with the ∼95% clinical CRS incidence for OKT3. Population-level CRS grade distributions across all three drugs are shown in Figure 5.

**Figure 4:**
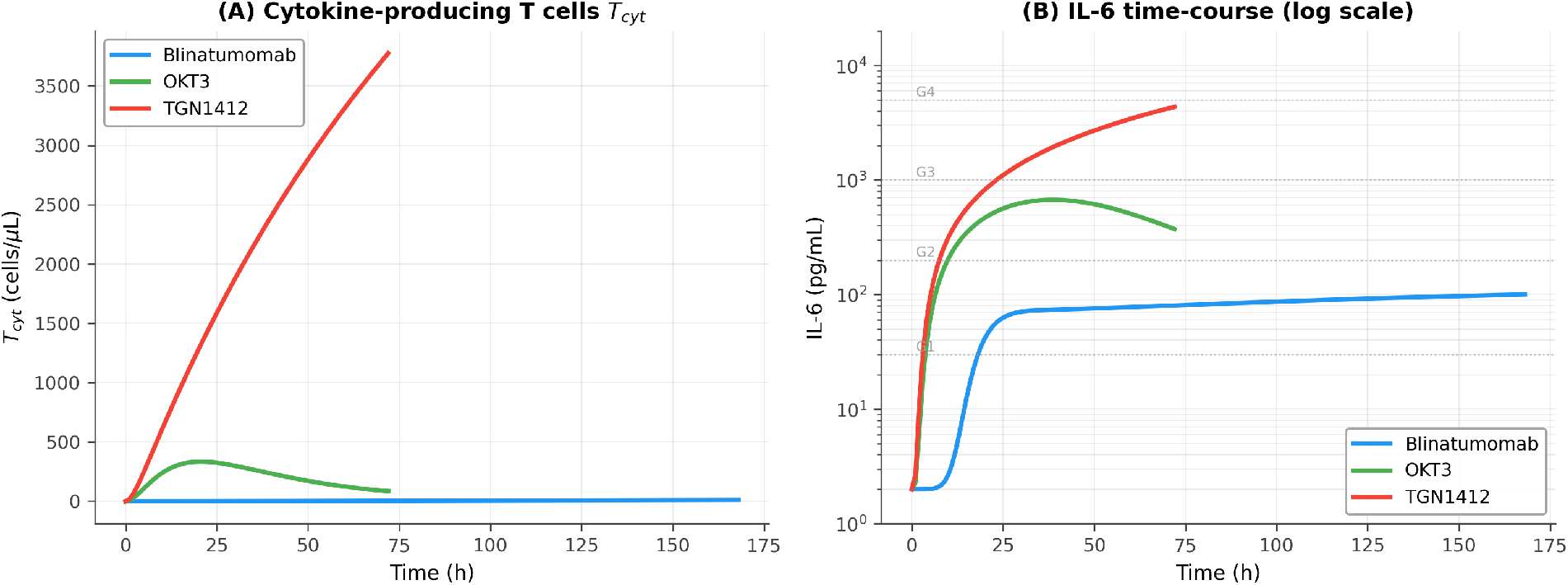
T cell mechanism determines CRS severity. (A) Cytokine-producing T cell (*T*_*cyt*_) dynamics for blinatumomab (blue), OKT3 (green), and TGN1412 (red). OKT3 shows peak-and-decline (AICD), while TGN1412 shows sustained expansion. (B) Resulting IL-6 dynamics (log scale). T cell expansion drives macrophage activation and IL-6 amplification through the shared cytokine network.

**Figure 5:**
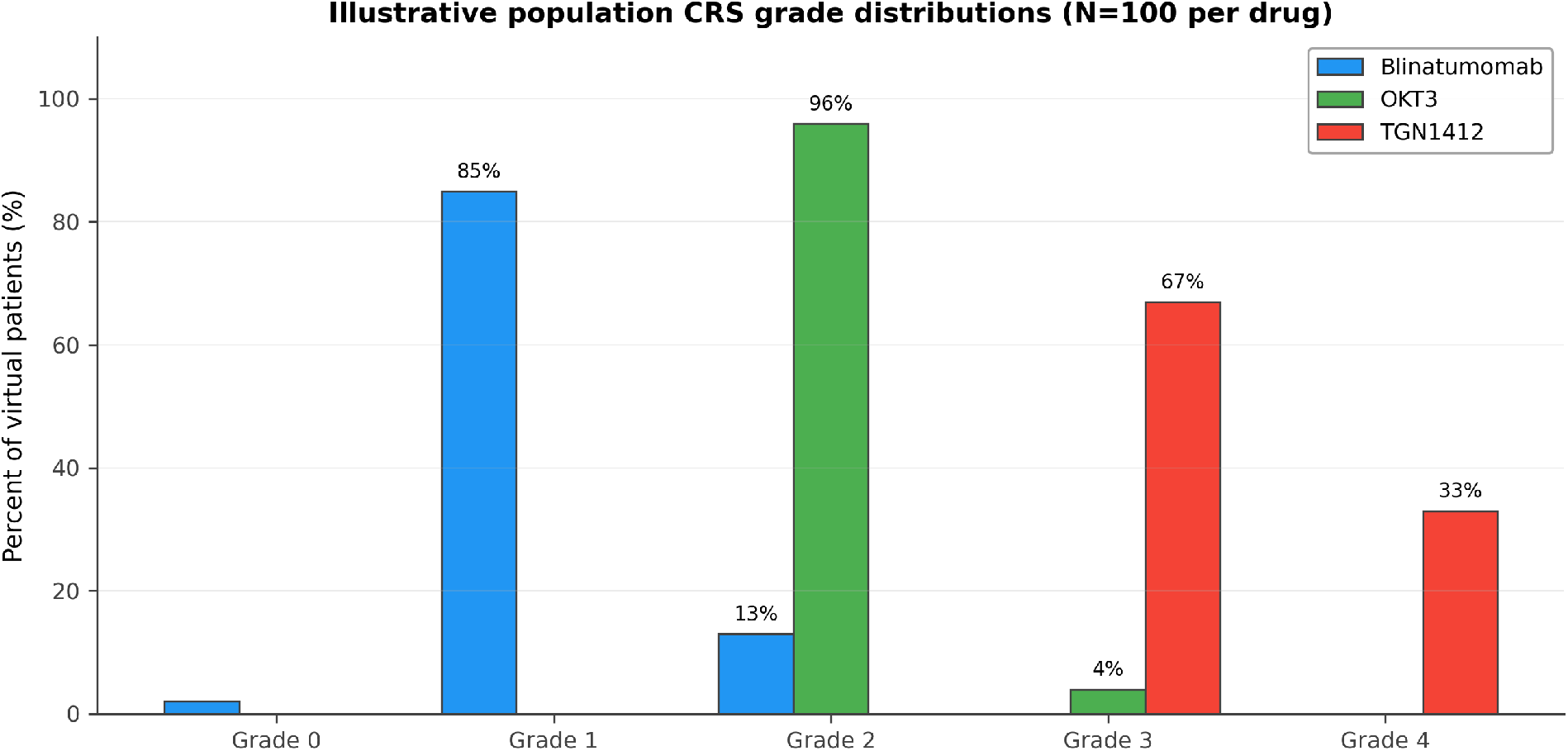
Illustrative CRS grade distributions across virtual populations (*N* = 100 per drug). Inter-individual variability was sampled from log-normal distributions (CV = 25-30%) on T cell activation parameters (*k*_*act*_, *k*_*prolif*_, *k*_*exhaust*_, *EC*_50,*act*_, *T*_*act,max*_), macrophage pool size, and body weight (see Methods for assumptions and limitations). Blinatumomab: predominantly Grade 1 (∼85%) with ∼13% Grade 2 and ∼2% sub-clinical. OKT3: ∼96% Grade 2 with ∼4% Grade 3, reflecting the AICD ceiling on CRS severity. TGN1412: split between Grade 3 (∼67%) and Grade 4 (∼ 33%), qualitatively consistent with clinical Grade 3-4 in all six Suntharalingam 2006 subjects. These distributions reflect sensitivity to assumed parameter variability rather than formally calibrated population predictions.

### 3.4 Parameter swap test

The parameter swap test was designed to distinguish between mechanistic prediction and parameter fitting (Table 6).

**Table 6.** Parameter swap test: exchanging *k*_*exhaust*_ between OKT3 and TGN1412.

| Condition | $k_{exhaust}$ | CRS Grade | Peak IL-6 | Interpretation |
| --- | --- | --- | --- | --- |
| OKT3 real (AICD) | 0.08 | 2 | 671 | Correct |
| OKT3 without AICD | 0.012 | 3 | 1,554 | Becomes severe (G2→G3) |
| TGN1412 real (expansion) | 0.012 | 3 | 4,349 | Correct |
| TGN1412 with forced AICD | 0.08 | 3 | 2,358 | IL-6 drops 46% (G3, attenuated) |

Swapping the single parameter $k_{exhaust}$ (T cell exhaustion/AICD rate) between OKT3 and TGN1412 reverses the direction of CRS severity: OKT3 crosses from Grade 2 to Grade 3 when AICD is removed, and TGN1412 peak IL-6 drops by 46% when AICD is imposed (although both TGN1412 conditions remain Grade 3 in the current calibration). The effect of the swap is in the expected direction for both drugs, supporting the interpretation that the model captures a real biological distinction (activation-induced cell death versus sustained expansion) rather than relying on arbitrary parameter tuning.

Removing AICD from OKT3 (*k*_*exhaust*_ : 0.08 *→* 0.012) increased severity from Grade 2 to Grade 3 (IL-6: 671 *→* 1,554 pg/mL). Forcing AICD on TGN1412 (*k*_*exhaust*_ : 0.012 *→* 0.08) dropped peak IL-6 by 46% (4,349 *→* 2,358 pg/mL), although both the real and AICD-forced TGN1412 conditions remain Grade 3 in the current calibration. The direction of change is correct in both swaps: the single mechanistically meaningful parameter controls whether a drug sits in the low-to-moderate or high-severity regime, confirming that the T cell mechanism, not arbitrary parameter tuning, drives the CRS prediction (Figure 6).

**Figure 6:**
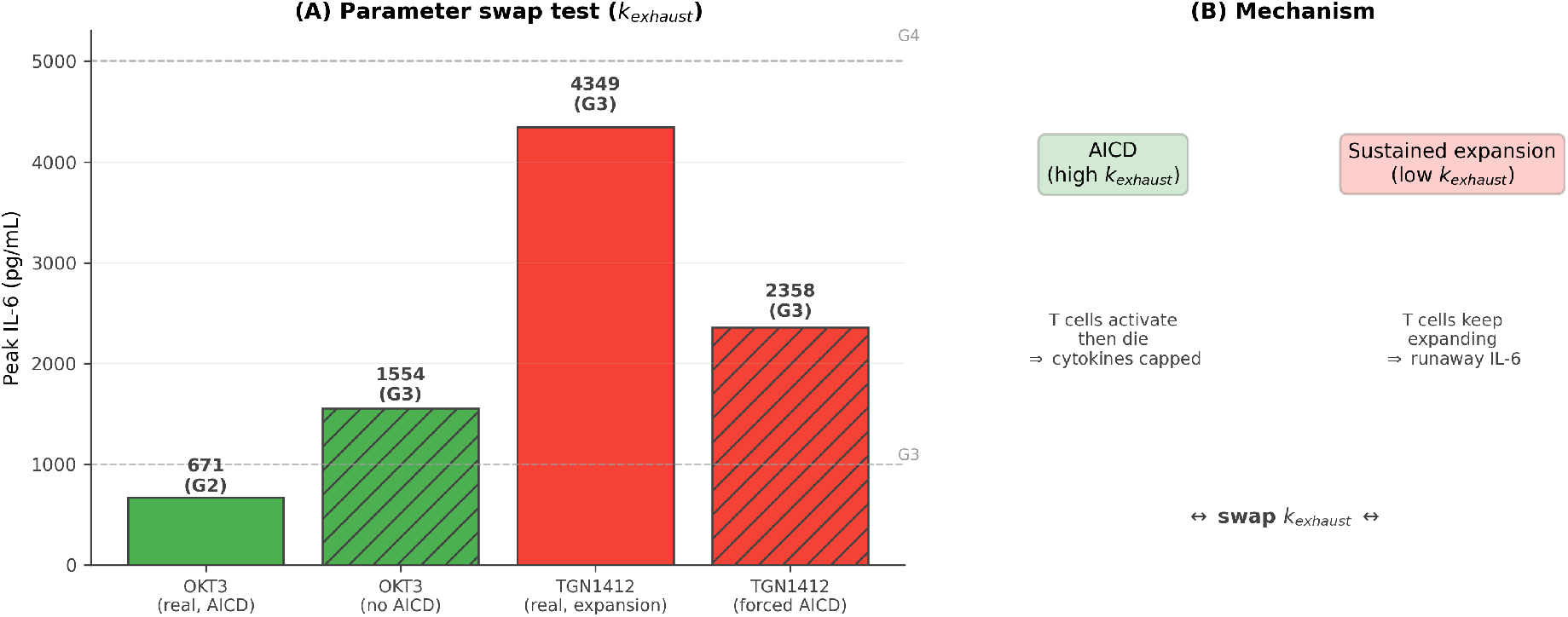
Parameter swap alters CRS severity in the expected direction. (A) Peak IL-6 under four conditions: OKT3 with real AICD (*k*_*exhaust*_ = 0.08, Grade 2, 671 pg/mL), OKT3 without AICD (*k*_*exhaust*_ = 0.012, Grade 3, 1,554 pg/mL), TGN1412 with real expansion (*k*_*exhaust*_ = 0.012, Grade 3, 4,349 pg/mL), TGN1412 with forced AICD (*k*_*exhaust*_ = 0.08, Grade 3, 2,358 pg/mL). Hatched bars indicate swapped parameters. (B) Mechanistic schematic: AICD (T cells activate then die) limits cytokine production, while sustained expansion drives runaway IL-6. The swap moves OKT3 from Grade 2 to Grade 3 and drops TGN1412 peak IL-6 by 46%, supporting the interpretation that *k*_*exhaust*_ is the mechanistically meaningful switch between sustained-expansion and AICD-capped CRS.

### 3.5 Tocilizumab rescue simulation

The locked model reproduced five clinically validated phenomena for tocilizumab rescue of TGN1412-severity CRS, without any changes to the shared cytokine network (Table 7, Figure 7).

**Table 7.** Tocilizumab rescue: five blind predictions matching clinical evidence.

| Model Prediction | Clinical Evidence | Reference |
| --- | --- | --- |
| IL-6 amplification loop broken (89% reduction in effective IL-6) | Tocilizumab response within hours | Le 2018 [13] |
| Paradoxical IL-6 rise (raw levels increase under IL-6R blockade) | Well-documented post-tocilizumab IL-6 elevation | Nishimoto 2008 [14] |
| TNF-α/IFN-γ completely unaffected by tocilizumab | IL-6R blockade does not suppress other cytokines | Mechanism of action |
| Tocilizumab alone insufficient for TGN1412-severity CRS (TNF-α/IFN-γ inflammation persists) | <60% response rate at Grade 4 CRS; steroids needed | Le 2018 [13], ASTCT guidelines [12] |
| Critical intervention window (~24 h to prevent Grade 3) | Early tocilizumab more effective (>85% at Grade 2) | Le 2018 [13] |

The key finding was the dissociation between IL-6 signaling and other inflammatory pathways. Post-tocilizumab, raw IL-6 levels rose paradoxically (receptor-mediated clearance blocked, contributing ∼70% of total IL-6 clearance), but IL-6 *signaling* was zero (STAT3 feedback eliminated). Meanwhile, TNF-*α* and IFN-*γ* trajectories were *identical* with and without tocilizumab (peak TNF-*α* = 558 pg/mL and peak IFN-*γ* = 1,674 pg/mL regardless of treatment). This emerged from the model structure: T cell activation and IFN-*γ*/TNF-*α* production are IL-6-independent pathways; only the IL-6 autocrine STAT3 loop is disrupted by IL-6R blockade.

This structural prediction matches clinical practice: ASTCT guidelines [12] recommend tocilizumab for Grade 2 CRS but add corticosteroids for refractory or Grade *≥*3 CRS, precisely because tocilizumab alone cannot control the non-IL-6 inflammatory component.

**Figure 7:**
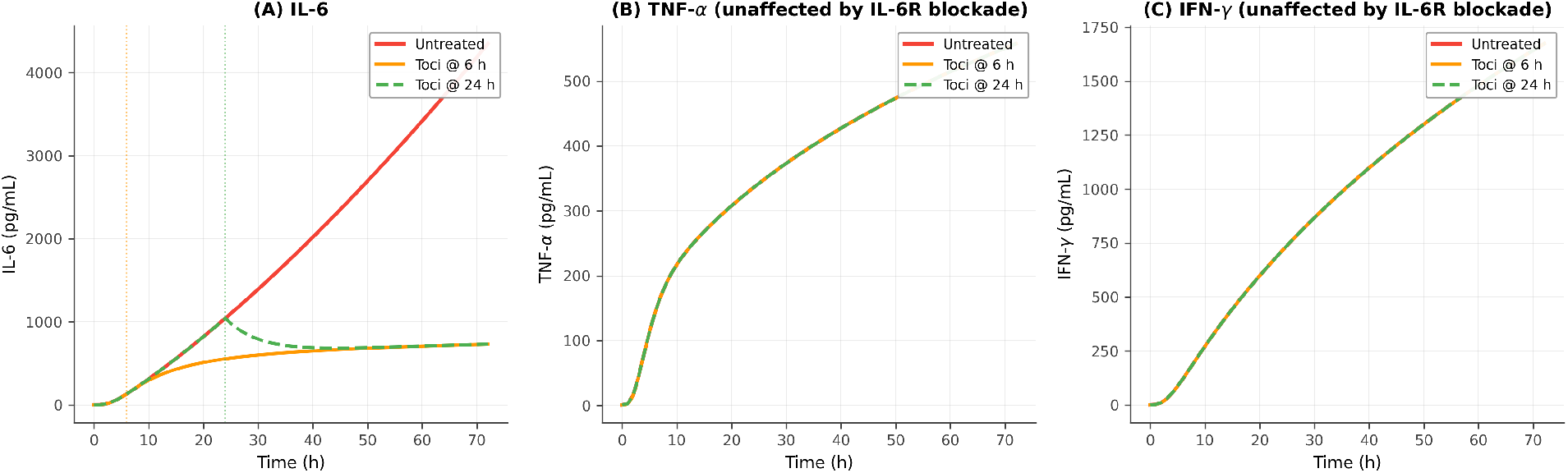
Tocilizumab rescue of TGN1412-severity CRS. (A) IL-6 time-course: untreated (red), tocilizumab at 6 h (orange), tocilizumab at 24 h (green dashed). IL-6 amplification loop is broken by IL-6R blockade. (B) TNF-*α* and (C) IFN-*γ* are completely unaffected by tocilizumab; trajectories are identical regardless of treatment, reflecting the IL-6-independent nature of T cell activation and macrophage inflammatory signaling. This emergent structural prediction matches ASTCT clinical guidelines: tocilizumab alone is insufficient for severe CRS.

### 3.6 Dose-response analysis

Dose-response analysis revealed three qualitatively distinct curve shapes, all from the same locked cytokine network (Figure 8).

#### Blinatumomab: gradual

Peak IL-6 rose smoothly from 71 to 226 pg/mL across the dose range (0.001-0.1 mg). Grade 1 was predicted across 9 of 10 dose levels, with only the highest dose crossing marginally into Grade 2 (226 pg/mL at 0.1 mg). This gradual response reflects low macrophage activation (*f*_*mac*_ *≈* 0.3) keeping IL-6 feedback well below the bifurcation threshold.

#### TGN1412: switch-like

Grade 1 at doses *≤* 0.00066 mg/kg (peak IL-6 < 100 pg/mL), jumping to Grade 3 at 0.0017 mg/kg (peak IL-6 ∼2,200 pg/mL) and beyond. This sharp threshold behavior emerges from two features: (i) the Hill coefficient *n*_*H*_ = 3.0 (cooperative CD28 signaling), and (ii) the bifurcation-like STAT3 amplification regime: once macrophage activation crosses a threshold, the effective feedback rate approaches the degradation rate and IL-6 production becomes explosive. The switch-like dose-response is consistent with the TGN1412 disaster, where the tested dose was only ∼1,000-fold above the subsequently safe Theralizumab dose.

#### OKT3: plateau

Peak IL-6 ranged from 600 to 671 pg/mL across a 40-fold dose range (0.5-20 mg). All doses produced Grade 2 CRS. This plateau reflects the AICD mechanism: even at high OKT3 doses, T cells activate then die, creating a natural ceiling on cytokine production that cannot be overcome by increasing dose.

These three qualitatively different shapes (gradual, switch-like, and plateau) are not individually fit; they emerge from the locked model with no dose-specific tuning.

**Figure 8:**
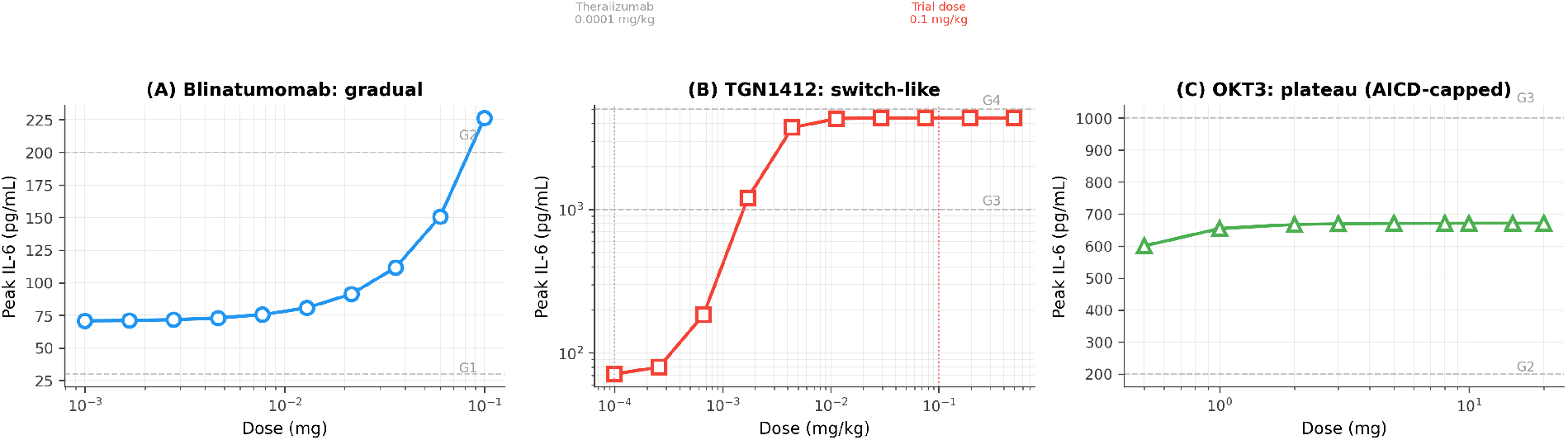
Three dose-response shapes from one locked model. (A) Blinatumomab: gradual IL-6 increase (71-226 pg/mL) across three orders of magnitude in dose; Grade 1 across nine of ten dose levels, with only the highest dose (0.1 mg) crossing marginally into Grade 2. (B) TGN1412: switch-like transition on log scale; Grade 1 below ∼0.001 mg/kg, Grade 3 above. Vertical dashed lines mark the Theralizumab safe dose (0.0001 mg/kg) and the TGN1412 trial dose (0.1 mg/kg). The sharp threshold reflects cooperative CD28 signaling (*n*_*H*_ = 3.0) combined with bifurcation-like STAT3 amplification regime. (C) OKT3: plateau from 600 to 671 pg/mL across a 40-fold dose range; AICD caps severity at Grade 2 regardless of dose. All panels use the same locked cytokine network parameters.

### 3.7 Local robustness across an accepted parameter ensemble

To test whether the cross-drug architectural transferability depends on the precise calibrated parameter values or holds across a wider feasible region around the calibrated point, a local ABC-style accepted parameter ensemble was constructed (Methods, Section 2.5). This is a local robustness analysis around the calibrated region, not a full Bayesian posterior, and the acceptance rule checks peak IL-6 only across the three antibody-based drugs (CAR-T is not included). From a uniform *±*30% prior around the calibrated values of the five free cytokine network parameters, 5,000 candidate parameter sets were sampled and each was evaluated.

**692 of 5**,**000 candidate parameter sets (13.8%) were accepted**, meaning that for each accepted set the model produced blinatumomab peak IL-6 within 50-200 pg/mL, TGN1412 peak IL-6 within 3,000-100,000 pg/mL, and OKT3 peak IL-6 within 200-2,000 pg/mL simultaneously. Of the 4,308 rejected candidates, 1,831 (36.6%) triggered IL-6 amplification runaway and 2,477 produced at least one drug outside the clinical range (TGN1412 was the most common single failure: 2,300 cases, almost always falling below 3,000 pg/mL). The 13.8% acceptance rate within a *±*30% local neighborhood indicates that the calibrated point is not an isolated solution but lies in a substantial feasible region of the cytokine network parameter space, characterized within this local neighborhood.

The accepted parameter distributions are tightly clustered around the calibrated values (Table 8; full distributional plots in Supplementary Figure S1). Median accepted values match the calibrated values to within 1% for all five parameters, with 5-95th percentile ranges spanning approximately *±*25% of the calibrated point. This indicates that the calibrated point sits at the center of the feasible region, not at its edge.

The cross-drug severity ordering is also robust across the accepted ensemble. Across all 692 accepted parameter sets, blinatumomab is predicted as Grade 1 in 100% of cases (median peak IL-6 = 101 pg/mL, 5-95th: 55-176 pg/mL), TGN1412 is predicted as Grade 3 in 28% and Grade 4 in 72% of cases (median peak IL-6 = 9,936 pg/mL, 5-95th: 3,220-71,677 pg/mL),and OKT3 is predicted as Grade 2 in 95% and Grade 3 in 5% of cases (median peak IL-6 = 642 pg/mL, 5-95th: 400-1,012 pg/mL). (The TGN1412 accepted-ensemble median of 9,936 pg/mL is higher than the single-patient baseline of 4,349 pg/mL because the acceptance range [3,000-100,000 pg/mL] is very broad: the sampling process retains parameter sets that shift TGN1412 IL-6 upward as well as downward, and the broad upper range means that the accepted distribution is right-skewed relative to the calibrated point.) The “Grade 3-4” TGN1412 predictions across the ensemble are consistent with the Suntharalingam 2006 NEJM clinical observation of “Grade 3-4 in all six volunteers,” and the “Grade 2-3” OKT3 predictions are consistent with the Chatenoud 1989 observation of Grade 2-3 first-dose reaction.

**Table 8.** Accepted parameter ensemble summary (5,000 samples, 692 accepted). Median and 5-95th percentile values across the accepted parameter sets, compared with the calibrated point.

| Parameter | Calibrated | Accepted median | 5% | 95% |
| --- | --- | --- | --- | --- |
| $k_{IL6,fb}$ ($\text{h}^{-1}$) | 0.528 | 0.524 | 0.388 | 0.660 |
| $mac_{pool}$ ($\text{cells}/\mu\text{L}$) | 465 | 466 | 348 | 587 |
| $k_{prod,IL6,mac}$ ($\text{pg}/\text{mL}/\text{h}/\text{cell}$) | 0.108 | 0.110 | 0.081 | 0.138 |
| $k_{deg,IL6}$ ($\text{h}^{-1}$) | 0.519 | 0.521 | 0.390 | 0.657 |
| $EC_{50,mac}$ ($\text{pg}/\text{mL}$) | 48.6 | 48.9 | 35.3 | 61.6 |

### 3.8 Stability map *k*_*IL*6,*fb*_ **vs** *k*_*deg,IL*6_

To visualize the structure of the feasible region, a 2D stability map was generated over the two parameters most directly involved in the IL-6 STAT3 amplification competition (Methods, Section 2.5). Across the 30 *×* 30 grid (900 parameter combinations), 122 (13.6%) produced all three drugs within their clinical IL-6 ranges, 351 (39.0%) triggered runaway amplification in at least one drug, 427 (47.4%) were feasible but produced at least one drug outside the clinical range, and **0 produced wrong cross-drug ordering** (Figure 9). The 13.6% all-in-range fraction matches the 13.8% acceptance rate from the independent ABC ensemble, providing convergent evidence for the size of the feasible region.

**Figure 9:**
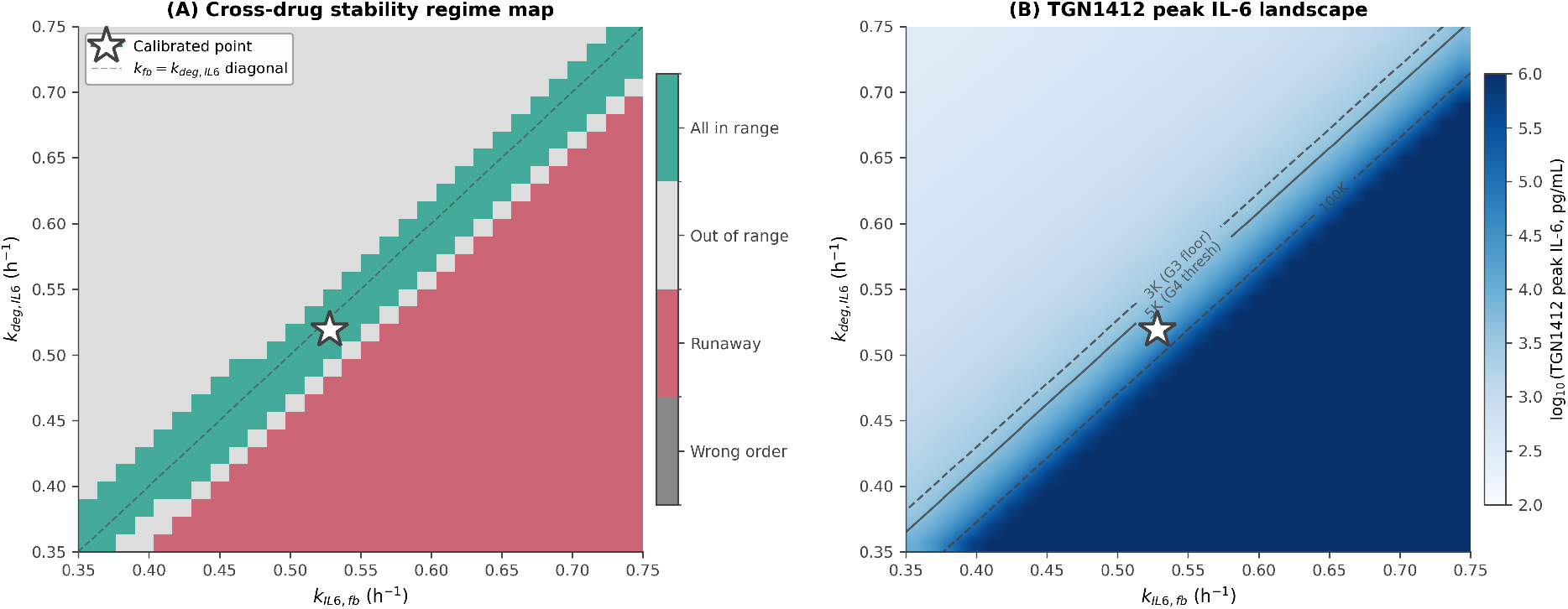
Cross-drug stability map for the two threshold-setting parameters. Colours use the Paul Tol muted qualitative palette in panel (A) and a single-hue blue sequential colourmap in panel (B); both schemes are colourblind-safe under deuteranopia, protanopia, and tritanopia. (A) Regime classification across a 30*×*30 grid of (*k*_*IL*6,*fb*_, *k*_*deg,IL*6_). Muted teal: all three antibody-based drugs predict peak IL-6 within their published clinical ranges. Light grey: feasible but at least one drug outside its clinical range. Muted rose-red: at least one drug undergoes IL-6 amplification runaway. Medium grey would indicate wrong cross-drug severity ordering (never observed in the data). Dashed line: theoretical bifurcation diagonal *k*_*IL*6,*fb*_ = *k*_*deg,IL*6_ (the boundary under maximal macrophage activation). White star: calibrated parameter point. Of 900 grid points, 122 (13.6%) fall in the muted-teal “all in range” region; 0 produced wrong cross-drug severity ordering. (B) Predicted TGN1412 peak IL-6 across the same grid (log scale, single-hue blue sequential colourmap; darker blue indicates higher peak IL-6), with dark-grey contour lines marking the 3,000 pg/mL Grade 3 floor, the 5,000 pg/mL Grade 4 threshold, and the 100,000 pg/mL upper clinical bound.

The stability map shows that the runaway region occupies the upper-left portion of the (*k*_*IL*6,*fb*_, *k*_*deg,IL*6_) plane (high feedback, low clearance), the safe-but-too-mild region occupies the lower-right (low feedback, high clearance), and the feasible region forms a band along the line *k*_*IL*6,*fb*_ *≈ k*_*deg,IL*6_ where the effective IL-6 amplification is poised at the boundary between self-limiting and runaway. The calibrated point sits inside this feasible band. This visual confirms the central mechanistic claim of the paper: cross-drug CRS severity separation is a property of the parameter region near the IL-6 STAT3 amplification boundary, not a property of any single tuned point.

### 3.9 Multi-cytokine acceptance test of the locked network

The local accepted parameter ensemble in Section 3.7 uses an IL-6-only acceptance rule. To test whether the locked cytokine network produces self-consistent multi-cytokine outputs, the same 5,000-sample local prior was re-evaluated against an extended acceptance rule that adds two further cytokines for TGN1412 (IFN-*γ* in 1,000-5,000 pg/mL and TNF-*α* in 1,000-3,500 pg/mL, both from Suntharalingam 2006 [4]) and two further cytokines for OKT3 (IFN-*γ* in 50-500 pg/mL from Ferran 1990 [7] and TNF-*α* in 100-800 pg/mL from Abramowicz 1989 [6] / Ferran 1990 [7]). Blinatumomab cytokines other than IL-6 are not consistently reported in Teachey 2013 case-series and were left out of the rule to keep it fair. The full acceptance rule therefore checks 7 cytokine-by-drug constraints simultaneously.

The honest result is informative on two levels. First, with all 7 constraints active, **0 of 5**,**000 samples were accepted**. Second, the dominant failure mode was **TGN1412 TNF-***α* (3,169 of 5,000 samples failed this single check), and the calibrated point itself produces TGN1412 TNF-*α* = 558 pg/mL, approximately 1.8-fold below the 1,000 pg/mL lower clinical bound from Suntharalingam 2006. This is the same TNF-*α* under-prediction documented in Limitations (Section 5.4, AFE = 3.35 against the geometric midpoint of the clinical range): the locked cytokine network calibrated on bispecific T cell engagers cannot reproduce the full magnitude of TGN1412 TNF-*α* within *±*30% of the calibrated parameters. The 0/5,000 result therefore confirms a known structural limitation of the network rather than indicating a new failure.

To characterize the robustness of the remaining cytokines, the same multi-cytokine ABC was re-run with the single failing constraint (TGN1412 TNF-*α*) excluded. Under this 6-constraint rule, **692 of 5**,**000 samples were accepted (13.8%), exactly matching the IL-6-only acceptance rate**. No additional samples were rejected by the IFN-*γ* checks for TGN1412 or OKT3, the TNF-*α* check for OKT3, or any combination of these. This means the IL-6 axis is the binding constraint on the locked network across this parameter neighborhood: any parameter set that produces blinatumomab, TGN1412, and OKT3 IL-6 within their clinical ranges automatically also produces TGN1412 IFN-*γ*, OKT3 IFN-*γ*, and OKT3 TNF-*α* within their clinical ranges. The locked network therefore produces self-consistent multi-cytokine outputs across the local feasible region for 6 of the 7 testable cytokine-by-drug constraints.

Reported transparently: (i) the full 7-constraint test yields 0/5,000 because of the known TGN1412 TNF-*α* structural failure; (ii) the 6-constraint conditional test yields 692/5,000 = 13.8%, identical to the IL-6-only result; (iii) the locked network passes 6 of 7 cytokine constraints throughout the entire feasible region but fails the 7th (TGN1412 TNF-*α*) at every point including the calibrated point.

### 3.10 Practical identifiability of the threshold-setting parameters

To assess whether the two parameters most directly involved in the IL-6 STAT3 amplification competition (*k*_*IL*6,*fb*_ and *k*_*deg,IL*6_) are practically identifiable individually, profile likelihood analyses were performed (Methods). For each target parameter, the parameter was fixed at a 21-point grid spanning *±*30% of the calibrated value, and the other four free parameters were re-optimized at each fixed value using the same constrained calibration objective (Section 2.5). Because the calibration objective is range-based (zero whenever all three antibody-based drugs lie within their published clinical IL-6 ranges, 10^6^ hard-rejection if any drug undergoes IL-6 runaway), the resulting profile is **not** a classical smooth likelihood curve but a **feasibility profile**: at each grid value the re-optimized four-parameter fit either reaches *J* (*θ*) = 0 (fully feasible under the joint cross-drug constraints) or is rejected. Feasible and rejected grid points therefore separate cleanly in Supplementary Figure S2, and the informative quantity is the *interval of feasibility* along each parameter axis rather than a curvature at an optimum.

The two profiles show **complementary asymmetric structure** (Supplementary Figure S2). The *k*_*IL*6,*fb*_ profile reaches *J* = 0 for the contiguous interval [0.370, 0.591] h^*−*1^, then jumps abruptly into the runaway-rejected regime for *k*_*IL*6,*fb*_ 0.607 h^*−*1^ (sharp upper boundary). The *k*_*deg,IL*6_ profile shows the opposite asymmetry: it is rejected for *k*_*deg,IL*6_ 0.472 h^*−*1^ (low clearance triggers runaway when the other parameters are re-optimized to match the IL-6 target), and reaches *J* = 0 for [0.488, 0.675] h^*−*1^ (soft upper boundary). The calibrated values (*k*_*IL*6,*fb*_ = 0.528, *k*_*deg,IL*6_ = 0.519) sit comfortably inside both feasible intervals, away from both runaway boundaries.

The biological interpretation is that **neither parameter is practically identifiable individually** within this range; each can vary by tens of percent and the other free parameters compensate to keep the calibration objective at zero. However, the two parameters are jointly constrained: *k*_*IL*6,*fb*_ has a sharp upper boundary at ∼0.60 h^*−*1^ while *k*_*deg,IL*6_ has a sharp lower boundary at ∼0.48 h^*−*1^, and both boundaries correspond to the same physical event (the IL-6 STAT3 feedback effective rate *k*_*IL*6,*fb*_ · *f*_*mac*_ exceeding the IL-6 clearance rate *k*_*deg,IL*6_, triggering runaway amplification). The two parameters are therefore practically identifiable as a ratio (or equivalently, as a difference *k*_*IL*6,*fb*_ *− k*_*deg,IL*6_ under maximal macrophage activation), not individually. This is consistent with the bifurcation interpretation already presented in the dose-response analysis and the stability map: the cross-drug severity architecture depends on the position of *k*_*IL*6,*fb*_ · *f*_*mac*_ relative to *k*_*deg,IL*6_, which is a ratio constraint, not a constraint on either parameter alone.

#### Interpreting the robustness numbers

The accepted fractions reported in Sections 3.7-3.10 should not be read as success rates in a single-drug fit. Every accepted parameter set had to satisfy *all* predefined clinical IL-6 range constraints *simultaneously* across the three antibody-based drugs (blinatumomab, TGN1412, OKT3) in a nonlinear near-threshold system; for the multi-cytokine variant, accepted sets also had to satisfy additional IFN-*γ* and TNF-*α* constraints. Under these joint constraints, the key question is not whether the percentage is large, but whether a non-trivial feasible region exists at all, and whether the calibrated solution lies inside that region.

The 692 of 5,000 accepted parameter sets (13.8%) in the local ABC ensemble, the 122 of 900 all-in-range grid points (13.6%) in the 2D stability map, and the 6 of 7 jointly satisfied cytokine constraints in the multi-cytokine acceptance test indicate that the calibrated solution is **not an isolated tuned point**: it lies within a finite, non-trivial local region of parameter space consistent with the observed cross-drug IL-6 behaviour. The two acceptance fractions (13.8% and 13.6%) agree closely despite coming from two independent methods (random local sampling in a 5-parameter neighbourhood versus a deterministic grid in a 2-parameter threshold plane), providing convergent evidence for the size of the feasible region.

The strongest single result from these analyses is that **0 of 900 stability-map grid points produced a wrong cross-drug severity ordering**, i.e., nowhere in the explored (*k*_*IL*6,*fb*_, *k*_*deg,IL*6_) plane was blinatumomab predicted more severe than OKT3, or OKT3 more severe than TGN1412. The severity ranking blinatumomab < OKT3 < TGN1412 is therefore *structurally preserved* across the entire explored threshold-parameter neighbourhood. This is stronger than the accepted fraction itself, because it shows that the relative severity ordering is a property of the architecture rather than of any specific calibrated point. The profile likelihood analysis is consistent with this interpretation: the two threshold-setting parameters are practically identifiable as a ratio (encoding the IL-6 amplification boundary) rather than individually, with the calibrated solution sitting comfortably away from the runaway edge in both profiles.

### 3.11 Consolidated validation summary for the antibody-based CRS claim

The locked cytokine network was subjected to a comprehensive set of complementary tests across the three antibody-based drug classes, including calibration on blinatumomab, structural refinement on TGN1412, blind prediction on OKT3 and tocilizumab rescue, mechanistic parameter swap, dose-response sweeps, and four robustness analyses (local accepted ensemble, stability map, multi-cytokine acceptance, and profile likelihood).

Table 9 summarizes the full set of validation tests, including their setup and outcome status. Table 10 reports the quantitative cytokine peak comparisons against published clinical ranges using two acceptance criteria: (i) whether each predicted peak falls within the reported clinical range, and (ii) the absolute fold-error relative to the geometric midpoint of the range.

**Table 9.** Consolidated validation of the cross-drug CRS transferability claim across the three antibody-based drug classes.

| Test | Setup / target | Result | Status |
| --- | --- | --- | --- |
| Three-drug CRS (blinatumomab) | Calibration on blinatumomab IL-6 range | Predicted Grade 1 (peak IL-6 = 100 pg/mL) | Calibration |
| Three-drug CRS (TGN1412) | Structural refinement on TGN1412 IL-6 range | Predicted Grade 3 (peak IL-6 = 4,349 pg/mL, within clinical 3-4) | Development |
| Three-drug CRS (OKT3) | 8 qualitative clinical plausibility checks | 8 / 8 pass | **Blind** |
| Quantitative cytokines (OKT3) | IL-6, TNF-α, IFN-γ vs. AFE $\leq 2$ | 3 / 3 pass | **Blind** |
| Tocilizumab rescue | 5 clinically validated phenomena against TGN1412-severity CRS | 5 / 5 match | **Blind** |
| Dose-response | Three drug classes, no dose-specific tuning | Gradual / switch-like / plateau shapes | **Blind** |
| Parameter swap test | $k_{exhaust}$ swap between OKT3 and TGN1412 | OKT3 G2→G3, TGN1412 IL-6 drops 46% | Mechanistic |
| Local accepted ensemble | 5,000 samples, $\pm 30\%$ local prior, IL-6-only acceptance on 3 drugs | 692 / 5,000 accepted (13.8%) | Robustness |
| Stability map | 900-point 2D grid on ($k_{IL6,fb}$, $k_{deg,IL6}$) | 0 / 900 wrong-order; 122 / 900 all-in-range | Robustness |
| Multi-cytokine acceptance | 7 cytokine-by-drug clinical-range constraints | 6 / 7 satisfied (TGN1412 TNF-α fails structurally) | Robustness |
| Profile likelihood | $k_{IL6,fb}$, $k_{deg,IL6}$ individual profiles | Complementary asymmetric; ratio-type practical identifiability | Identifiability |

#### Exploratory and supporting-module results in the Supplementary

An exploratory post hoc transfer test of the locked downstream cytokine network to tisagenlecleucel CAR-T cell therapy (pediatric and adult cohorts from Teachey 2016 [3]), with all methods, parameter choices, table, figure, and discussion, is provided in Supplementary Section S2. Validation results for the three supporting adverse event modules (IgG depletion, neutropenia, hepatotoxicity) are provided in Supplementary Sections S3-S5. Neither block is part of the main cross-drug CRS transferability claim of this paper.

**Table 10.** Quantitative comparison of predicted vs. observed cytokine peaks for the CRS module. Two acceptance criteria are applied: (i) whether the predicted peak falls within the reported clinical range, and (ii) absolute fold-error (AFE) relative to the geometric midpoint of the clinical range (AFE = max(pred/*G, G*/pred) where 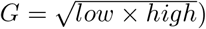, with AFE *≤* 2 as the acceptance threshold.

| Drug | Cytokine | Predicted<br>(pg/mL) | Clinical range<br>(pg/mL) | Geom. mid.<br>(pg/mL) | AFE | Status |
| --- | --- | --- | --- | --- | --- | --- |
| Blinatumomab | IL-6 | 100 | 50-200<sup>a</sup> | 100 | 1.00 | **Pass** |
| | TNF-α | 63 | - | - | - | No data |
| | IFN-γ | 9 | - | - | - | No data |
| TGN1412 | IL-6 | 4,349 | 3,000-100,000<sup>b</sup> | 17,321 | 3.98* | In range |
| | TNF-α | 558 | 1,000-3,500<sup>b</sup> | 1,871 | 3.35 | *Fail* |
| | IFN-γ | 1,674 | 1,000-5,000<sup>b</sup> | 2,236 | 1.34 | **Pass** |
| OKT3 | IL-6 | 671 | 200-2,000<sup>c</sup> | 632 | 1.06 | **Pass** |
| | TNF-α | 174 | 100-800<sup>c,d</sup> | 283 | 1.63 | **Pass** |
| | IFN-γ | 207 | 50-500<sup>c,d</sup> | 158 | 1.31 | **Pass** |

<sup>a</sup>Teachey 2013 [2]. <sup>b</sup>Suntharalingam 2006 [4]. <sup>c</sup>Chatenoud 1989 [5]. <sup>d</sup>Abramowicz 1989 [6], Ferran 1990 [7]. Out of 7 quantifiable cytokine peaks (blinatumomab TNF-α and IFN- $\gamma$ are excluded because clinical measurements were not reported), 6/7 fall within the reported clinical range and 5/7 additionally pass the 2-fold AFE criterion. \*TGN1412 IL-6: the prediction of 4,349 pg/mL is within the very broad clinical range of 3,000-100,000 pg/mL (a >30-fold span) but sits near its lower bound; because the geometric midpoint of this asymmetric range is 17,321 pg/mL, the AFE against the midpoint is 3.98 despite the prediction being in range. TGN1412 TNF-α fails both acceptance criteria (1.8-fold below the lower clinical bound of 1,000 pg/mL), consistent with the systematic TNF-α underestimation discussed in Limitations.

## 4 Sensitivity Analysis

One-at-a-time (OAT) parameter perturbation (*±*50%) was performed across all modules. The normalized sensitivity coefficient was *S* = (Δ*Y/Y*_0_)*/*(Δ*p/p*_0_).

For the CRS model, the macrophage pool size (*mac*_*pool*_, *S* = +1.71) and macrophage-driven IL-6 production rate (*k*_*prod,IL*6,*mac*_, *S* = +1.52) were the dominant parameters, consistent with monocyte-derived IL-6 as the primary CRS driver [8]. The IL-6 STAT3 feedback gain (*k*_*fb*_) exhibited near-threshold behavior: at *±*20% perturbation *S* = +2.9, but at +30% the system transitioned to runaway amplification (peak IL-6 > 16,000 pg/mL). This indicates that the baseline parameterization lies near the boundary between self-limiting and catastrophic CRS. Figure 10 presents the CRS-module sensitivity ranking as a tornado plot.

Taken together, these results indicate that CRS severity in the model is controlled primarily by the strength of macrophage-mediated IL-6 production and the balance between IL-6 positive feedback and clearance. Parameters affecting upstream T cell activation modulate the magnitude of the upstream input, but the transition from controlled to severe CRS is governed by the downstream amplification boundary, consistent with the stability map and profile likelihood analyses (Sections 3.8 and 3.10).

Full OAT sensitivity tables for the three supporting adverse event modules (IgG depletion, neutropenia, hepatotoxicity) are reported in Supplementary Section S7.

**Figure 10:**
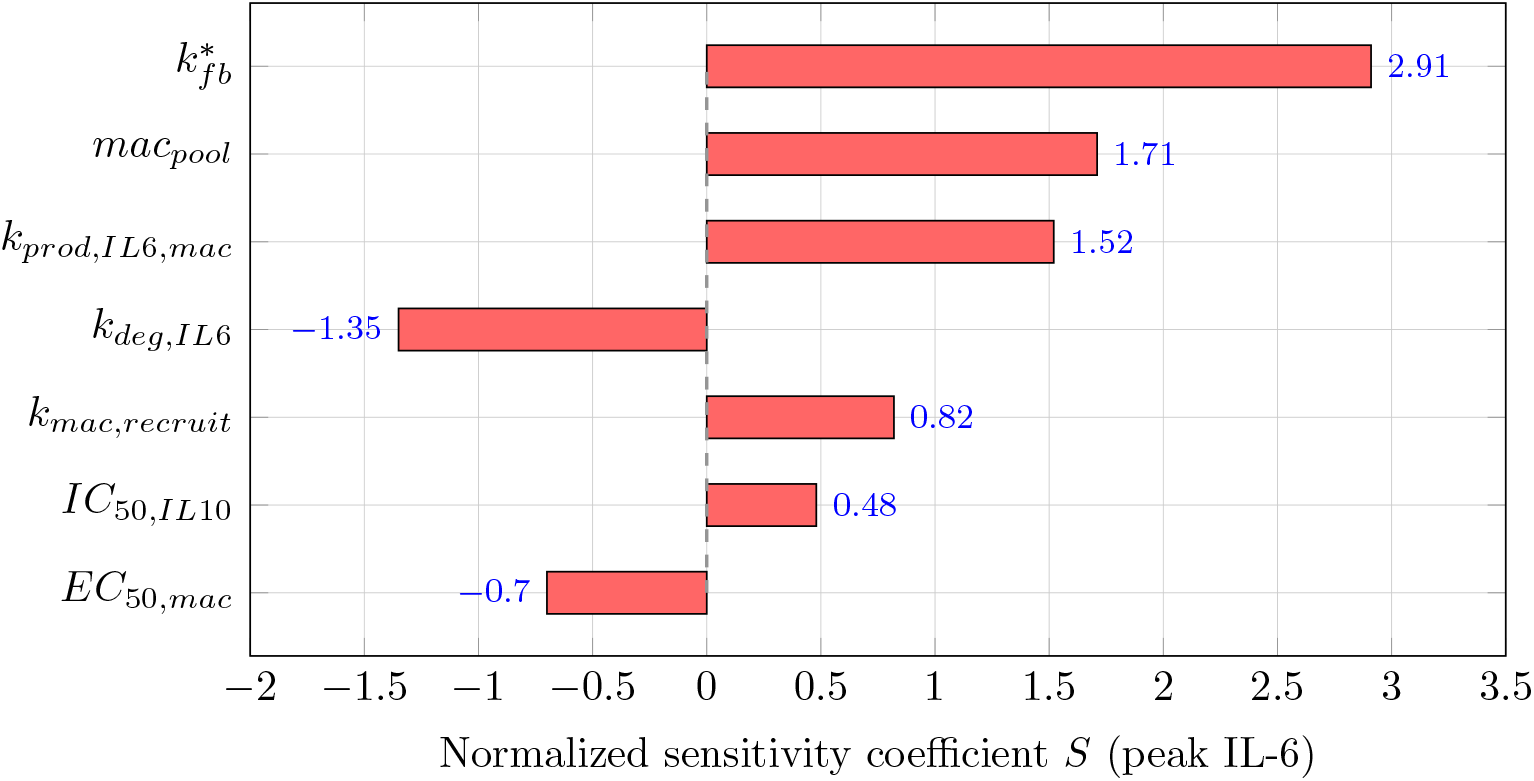
Tornado plot of one-at-a-time sensitivity coefficients for the CRS model. Each bar shows the normalized sensitivity *S* = (Δ*Y/Y*_0_)*/*(Δ*p/p*_0_) with peak IL-6 as the output, computed at *±*50% parameter perturbation from baseline (except 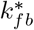, computed at *±*20% due to near-threshold behavior at larger perturbations). Parameters with |*S*| > 1 amplify the input change; parameters with |*S*| < 1 attenuate it. The dominance of *k*_*fb*_, *mac*_*pool*_, and *k*_*prod,IL*6,*mac*_ reflects the centrality of macrophage-derived IL-6 production and STAT3 positive feedback in driving CRS severity. One-at-a-time perturbation does not capture parameter interactions; variance-based global sensitivity methods (Morris, Sobol) are planned as a next step.

## 5 Discussion

This work demonstrates that a single 17-parameter cytokine network, calibrated on blinatumomab and structurally refined on TGN1412, retrospectively captures CRS severity for a third drug class (OKT3) and a therapeutic intervention (tocilizumab) without re-fitting the shared downstream network. The model also produces three qualitatively distinct dose-response shapes from the same locked parameters, and a single-parameter swap reverses the CRS severity ordering between drug classes.

### 5.1 Cross-drug validation

The three-drug result addresses a specific gap in the CRS modeling literature. Published CRS QSP models for CAR-T [3, 24] and bispecific antibodies [2, 15] are drug-class-specific. They capture cytokine dynamics for one agent but do not address whether the same network applies across mechanistically different drugs. The contribution here is therefore not the cytokine network per se, but the demonstration that one locked network generalizes across drug classes: a single cytokine network architecture, with only drug-specific T cell activation parameters varying, retrospectively captures severity differences spanning Grade 1 (blinatumomab) through Grade 2 (OKT3) to Grade 3 (TGN1412), corresponding to a ∼43-fold span in peak IL-6 (100 to 4,349 pg/mL).

The phrase “without re-fitting” refers specifically to the 17 shared downstream cytokine network parameters, which remained locked across all three drug classes and the tocilizumab intervention. Drug-specific upstream T cell activation parameters (6-7 per drug: *k*_*act*_, *k*_*exhaust*_, *k*_*prolif*_, *EC*_50,*act*_, *n*_*H*_, *T*_*act,max*_, *k*_*diff*_) were set independently for each agent. A subset of these parameters has direct literature support (e.g., *n*_*H*_ = 3.0 for TGN1412 from CD28 cooperative signaling [10], *k*_*exhaust*_ for OKT3 reflecting AICD kinetics from Chatenoud [5], IgG4 clearance range from Dirks 2010 [17]). Others reflect values drawn from QSP T cell engager models [22], assumed upper bounds (e.g., *T*_*act,max*_), or simplified PK representations (e.g., one-compartment approximations for OKT3 and blinatumomab). Full parameter provenance is tabulated in Table 1 footnotes, including honest acknowledgment of assumed values. The cross-drug claim therefore rests on architectural transferability of the shared CRS amplification network under drug-specific upstream dynamics, not on the absence of any drug-specific parameterization. The parameter swap test provides additional evidence that severity ordering follows from a mechanistic distinction (AICD vs. sustained expansion) rather than from the specific numerical values of the drug-specific parameters.

The mechanistic basis is the interaction between drug-specific T cell dynamics and shared macrophage-dependent IL-6 amplification. Blinatumomab activates a limited T cell subset, producing low IFN-*γ*/TNF-*α* that weakly activates macrophages (*f*_*mac*_ *≈* 0.3), keeping effective IL-6 feedback below the bifurcation threshold. TGN1412 causes massive polyclonal T cell expansion (no AICD), producing high IFN-*γ*/TNF-*α* that maximally activates and recruits macrophages (*f*_*mac*_ *≈* 0.99, 5-fold pool amplification), pushing effective IL-6 feedback near the critical degradation rate. OKT3 initially activates T cells like TGN1412 but AICD limits T cell numbers within 24 h, capping macrophage activation and preventing bifurcation-like STAT3 amplification regime.

The dose-response shapes provide independent evidence for this mechanism. The gradual blinatumomab response (IL-6 always below the amplification threshold), switch-like TGN1412 response (threshold crossing), and OKT3 plateau (AICD-capped) are not individually parameterized; they emerge from the interaction between drug-specific T cell dynamics and the shared nonlinear cytokine network. These qualitatively different shapes are much harder to dismiss as parameter fitting than point predictions at individual doses.

### 5.2 Structural necessity of the macrophage-gated feedback

A natural concern with any mechanistic model is whether the structural features are genuinely required to explain the observations or whether simpler alternatives would also work. Two structural arguments in the present work constrain this concern.

First, an ablated version of the model without macrophage-gated STAT3 feedback (constitutive feedback with *k*_*fb*_ = 0.4 h^*−*1^) was investigated during model development (Section 2.4, Step 2). This ablated model predicted TGN1412 CRS as Grade 2 (peak IL-6 = 711 pg/mL), drastically underestimating the clinical Grade 3 to 4. Increasing *k*_*fb*_ above 0.5 to match TGN1412 caused all drugs (including blinatumomab) to produce Grade 4 CRS, destroying cross-drug severity separation. Cross-drug separation therefore requires a drug-dependent effective feedback rate; a constitutive feedback cannot simultaneously capture blinatumomab Grade 1 to 2 and TGN1412 Grade 3 to 4 with any single value of *k*_*fb*_.

Second, the macrophage gating (*f*_*mac*_) and recruitment (*k*_*mac,recruit*_) terms are not freefloating mathematical devices; they encode a specific biological hypothesis consistent with the Norelli and colleagues finding that monocyte-derived IL-6 is the primary driver of CRS severity [8]. One could reasonably ask whether the second amplification layer (macrophage recruitment in addition to STAT3 feedback) is convenience-driven. Both layers have independent biological support: STAT3 positive feedback is a well-characterized IL-6 autocrine mechanism, and IFN-*γ*-driven monocyte recruitment during inflammatory storms is documented in TGN1412 pathology. Neither is introduced ad hoc to fit the data; both are observed phenomena whose mathematical representation is required to reproduce the clinical severity hierarchy.

These structural arguments do not substitute for formal identifiability analysis, but they establish that the central model features are necessary (without macrophage gating, cross-drug separation fails) and biologically grounded (both feedback layers have published experimental support).

### 5.3 The tocilizumab prediction

The tocilizumab rescue simulation was a particularly stringent test because it probed model *structure*, not parameters. The prediction that TNF-*α* and IFN-*γ* are completely unaffected by IL-6R blockade emerges from the model architecture: T cell activation and macrophage activation by IFN-*γ*/TNF-*α* are IL-6-independent pathways. Only the IL-6 autocrine STAT3 loop is disrupted. This structural prediction matches ASTCT clinical guidelines [12]: tocilizumab alone is effective for Grade 2 CRS but Grade *≥*3 requires adding corticosteroids, precisely because non-IL-6 inflammation persists.

### 5.4 Limitations

Several limitations must be acknowledged.

#### Validation scope

##### TGN1412 is not a blind prediction

The macrophage-gated STAT3 feed-back was discovered because the model failed on TGN1412. While the improvements are biologically justified [8], TGN1412 must be characterized as a model development case, not a validation.

##### Not prospectively validated

All three drugs are well-studied historical cases. The model cannot yet claim prospective predictive utility, only that “retrospectively captures CRS severity differences across mechanistically distinct drug classes, suggesting potential for prospective application pending further validation.”

#### Structural limitations

##### TGN1412 TNF-α is under-predicted

For TGN1412 the model under-predicts peak TNF-*α* (558 vs. clinical geometric midpoint 1,871 pg/mL, AFE = 3.35, and approximately 1.8-fold below the lower clinical bound of 1,000 pg/mL). This is the single failing cytokine constraint identified by the quantitative comparison (Table 10) and by the multi-cytokine acceptance test, which rejects TGN1412 TNF-*α* at every parameter set tested including the calibrated point. It is a structural limitation of the bispecific-calibrated cytokine network and suggests that the macrophage TNF-*α* production pathway should be made drugclass-aware in a future revision. Because the CRS grade in the present model is anchored to peak IL-6 rather than peak TNF-*α*, this discrepancy does not affect the cross-drug severity ordering. The exploratory CAR-T transfer test reported in Supplementary Section S2 shows a distinct, opposite-direction TNF-*α* failure for cell therapy contexts; taken together, these two failure modes point to drug-class-dependent macrophage TNF-*α* biology that a single locked network cannot capture and that is flagged as a priority for future refinement.

#### Parameterization and identifiability

##### Drug-specific T cell parameters are partially free

Each drug has 6-7 T cell parameters (*k*_*act*_, *k*_*exhaust*_, *k*_*prolif*_, *EC*_50,*act*_, *n*_*H*_, *T*_*act,max*_, *k*_*diff*_), giving 18-21 total degrees of freedom across the three drugs. A potential concern is that with this many parameters, matching three drug classes is unsurprising. Partial mitigation: (i) the parameters most closely tied to CRS severity differences (*n*_*H*_ for TGN1412 cooperativity and *k*_*exhaust*_ for OKT3 AICD) have direct literature support; (ii) the parameter swap test demonstrates that severity ordering reverses when a single mechanistically meaningful parameter is exchanged between drugs, arguing against arbitrary parameter tuning; (iii) dose-response shapes emerge from the locked model without dose-specific adjustment. Remaining free parameters (*k*_*act*_, *k*_*diff*_, *k*_*prolif*_, *T*_*act,max*_) reflect QSP T cell engager literature values [22] or physiological bounds. Formal parameter identifiability analysis is needed to quantify this concern rigorously.

##### Calibration: formal constrained range-based, with local-region characterization and partial practical identifiability assessment

Calibration was performed by a constrained optimization with a range-based blinatumomab IL-6 target and soft cross-drug architectural preservation penalties, on a subset (5 of 17) of cytokine network parameters selected by prior OAT sensitivity ranking (Methods, Section 2.5). The robustness of the calibrated point was characterized by a local accepted parameter ensemble (Section 3.7, 13.8% acceptance rate within *±*30% local prior on the IL-6 axis), a 2D *k*_*IL*6,*fb*_-vs-*k*_*deg,IL*6_ stability map (Section 3.8, 0% wrong-order regime), a multi-cytokine acceptance test (Section 3.9, 6 of 7 cytokine-by-drug constraints simultaneously satisfied across the entire feasible region), and a profile likelihood analysis of the two threshold-setting parameters (Section 3.10, complementary asymmetric profiles consistent with ratio-based identifiability).

These analyses have three remaining scope limits. First, the prior is local (*±*30% around calibrated values) and uniform, not a biologically informed prior over the broader plausible parameter space; conclusions about feasibility apply to this local neighborhood only. Second, the multi-cytokine acceptance rule fails on TGN1412 TNF-*α* at every parameter set tested (discussed above under structural limitations); this affects multi-cytokine robustness but not the IL-6-anchored cross-drug severity ordering. Third, the procedure involves only the three antibody-based drugs; CAR-T is never part of the calibration, the ensemble, the stability map, the multi-cytokine test, or the profile likelihood, and remains exclusively in the post hoc exploratory transfer test reported in Supplementary Section S2. The present analyses provide **practical identifiability** (via profile likelihood and accepted ensemble) but not **full global identifiability** in the strict mathematical sense, which would require a biologically informed prior, a Bayesian or rank-based identifiability framework, and variance-based global sensitivity (Morris, Sobol). Users evaluating the model for regulatory or translational deployment should treat the present parameterization as one feasible configuration in a 13-14% slice of the local *±*30% parameter neighborhood that is jointly constrained by 6 of 7 cytokine targets with the seventh (TGN1412 TNF-*α*) being a documented structural limit.

### 5.5 Future directions

A proper CAR-T CRS model, including antigen-dependent engagement, tumor burden dynamics as an off-switch on CAR-T-driven cytokine release, and age- or disease-burden-dependent upstream parameters, is a key next direction, as the exploratory transfer test reported in Supplementary Section S2 identifies exactly these as the missing biology.

PK refinement to a two-compartment model with rapid distribution is a near-term priority and would improve the predicted CRS onset timing (currently 3-4 h vs. clinical 60-90 min for TGN1412 and 30-60 min for OKT3) without altering steady-state CRS severity predictions. Drug-specific PK parameter updates (blinatumomab *V*_*central*_ = 5.98 L per Clements 2020 [16] vs. current 3.5 L; OKT3 CL ∼0.13 L/h per Hooks 1991 [18] vs. current 0.02 L/h) would be incorporated in the same refinement.

For sensitivity and identifiability, Morris elementary-effects screening, computationally inexpensive at *∼k* · (*p* + 1) runs for *p* parameters, is the recommended immediate next step over the OAT analysis presented here. Variance-based global methods (Sobol indices) and sparse polynomial chaos expansions would extend this to a full identifiability assessment with parameter interactions and correlations explicitly characterized. Independently, formal parameter estimation and prospective validation against dose-escalation data from novel T cell engagers are also planned.

This work positions cytokine release syndrome modeling within a model-informed drug development (MIDD) framework, where exposure-driven immune activation propagates through a shared amplification network to determine clinical safety outcomes. The recently issued ICH M15 guideline on model-informed drug development [23] formally recognizes QSP models, including mechanistic safety models, as valid evidence for regulatory decision-making. The cross-drug architectural transferability demonstrated here, in which only drug-specific T cell activation parameters need specification for novel agents, aligns with the ICH M15 vision of platform models that support first-in-human dose selection across therapeutic programs. Virtual patient generation using the locked cytokine network could further support in silico clinical trial design for CRS risk stratification, an area of growing regulatory interest. Integration of machine learning surrogate approaches with the mechanistic framework, for example training gradient-boosted or neural network emulators on simulated virtual populations, could accelerate sensitivity analysis and enable rapid patient-specific CRS risk prediction without sacrificing mechanistic interpretability.

Taken together, the central contribution of this work is architectural transferability of a shared CRS amplification network within the antibody-based T cell engager class, with drug-specific T cell activation parameters as the only inputs that vary. This is an in silico mechanistic proof-of-concept, not definitive prospective clinical deployment; the latter requires the prospective validation, global identifiability assessment, and cell-therapy extension outlined above.

## Study Highlights

### What is the current knowledge on the topic?

Cytokine release syndrome (CRS) models exist for individual drug classes (CAR-T, bispecifics) but no single mechanistic cytokine network has been shown to capture CRS severity differences across mechanistically distinct drug classes without re-fitting the shared downstream network.

### What question did this study address?

Can a single locked cytokine amplification network, calibrated on one drug class and structurally refined on a second, reproduce CRS severity, dose-response shape, and therapeutic rescue for a third drug class and a rescue intervention, without re-fitting the shared downstream network?

### What does this study add to our knowledge?

A 17-parameter cytokine amplification network with macrophage-gated STAT3 feedback retrospectively captures CRS severity across three antibody-based drug classes (blinatumomab calibration, TGN1412 development case, OKT3 blind prediction with 3/3 cytokine peaks within clinical ranges and 8/8 plausibility checks), with the cross-drug severity ordering preserved across a 13.8% local feasible region characterized by an accepted parameter ensemble and a 2D stability map (0/900 wrong-order). Profile likelihood and multi-cytokine acceptance analyses show that the IL-6 axis is the binding constraint and that 6 of 7 published cytokine ranges are jointly satisfied across the feasible region. The same locked model predicts three distinct dose-response shapes and reproduces five clinically validated tocilizumab rescue phenomena. An exploratory post hoc transfer test of the locked downstream network to tisagenlecleucel CAR-T cell therapy is provided in the Supplementary Material and is not part of the main antibody-based claim.

### How might this change drug discovery, development, and/or therapeutics?

The cross-drug generalizability suggests this cytokine network architecture could support prospective CRS risk stratification for novel immunomodulatory agents, where only drug-specific T cell activation parameters need to be specified

## Data and Code Availability

All validation data were extracted from published sources cited in the text and are included in the locally-stored package as digitized datasets. The locked 17-parameter cytokine network listed in Table 2 corresponds to a formally calibrated parameter file used unchanged for all blind testing reported in this work. Full source code, the locked parameter file, and reproducibility scripts will be released in a public repository upon journal publication. Reasonable requests for code or data prior to publication can be addressed to the corresponding author.

