## Supplementary Material for "A PK-Driven Quantitative Systems Pharmacology Model Predicts Cytokine Release Syndrome Severity Across T Cell-Activating Therapies via a Locked Cytokine Amplification Network"

Hajar Besbassi*[1,2,3]

[1]Antwerp Centre for Translational Immunology and Virology (ACTIV), University of Antwerp, Antwerp, Belgium
[2]Centre for Health Economics Research and Modelling Infectious Diseases (CHERMID), University of Antwerp, Antwerp, Belgium
[3]Vaccine & Infectious Disease Institute (VAXINFECTIO), University of Antwerp, Antwerp, Belgium

This supplementary material supports the cross-drug CRS analysis presented in the main paper. It provides (i) the complete ODE system of the CRS model (Section S1); (ii) an exploratory post hoc transfer test of the locked downstream cytokine network to tisagenlecleucel CAR-T cell therapy (Section S2); (iii) the local accepted parameter ensemble distributional plots referenced in the main text (Section S3, Figure S1); (iv) the constrained feasibility profile analysis for the two threshold-setting parameters (Section S4, Figure S2); (v) the virtual population generator (Section S5), full one-at-a-time sensitivity analysis (Section S6), and parameter calibration procedure (Section S7). The exploratory CAR-T extension in Section S2 is explicitly not a blind validation and is reported here rather than in the main paper because CAR-T-specific upstream parameters were formulated with knowledge of the Teachey 2016 cytokine data.

#### S1 CRS Model: Complete ODE System

The CRS model in the main manuscript is described via individual mechanistic equations. This section provides the full 11-state ODE system in explicit form for reproducibility.

##### S1.1 State variables

The CRS model has 11 state variables: drug concentration $C_{drug}$ (nM); four T cell compartments $T_{rest}$, $T_{act}$, $T_{cyt}$, $T_{exh}$ (cells/$\mu$L); and five cytokine concentrations $[IFN\gamma]$, $[TNF\alpha]$, $[IL6]$, $[IL1\beta]$, $[IL10]$ (pg/mL). Macrophage activation $f_{mac}$ is an auxiliary quasi-static function, not a dynamical state.

#### S1.2 Auxiliary functions

T cell activation Hill function:

$$f_{act}(C_{drug}) = \frac{C_{drug}^{n_H}}{EC_{50,act}^{n_H} + C_{drug}^{n_H}} \tag{S1}$$

Macrophage activation fraction:

$$f_{mac} = \frac{[IFN\gamma] + [TNF\alpha]}{EC_{50,mac} + [IFN\gamma] + [TNF\alpha]} \tag{S2}$$

Effective macrophage pool with IFN-$\gamma$/TNF-$\alpha$-driven recruitment:

$$mac_{pool,eff} = mac_{pool} \cdot \left(1 + k_{mac,recruit} \cdot \frac{[IFN\gamma] + [TNF\alpha]}{K_{m,recruit} + [IFN\gamma] + [TNF\alpha]}\right) \tag{S3}$$

IL-10 anti-inflammatory suppression:

$$f_{IL10} = \frac{1}{1 + [IL10]/IC_{50,IL10}} \tag{S4}$$

#### S1.3 Drug pharmacokinetics

$$\frac{dC_{drug}}{dt} = \frac{R_{inf}(t)}{V_{central}} - \frac{CL}{V_{central}} \cdot C_{drug} \tag{S5}$$

#### S1.4 T cell state dynamics

$$\frac{dT_{rest}}{dt} = -k_{act} \cdot f_{act} \cdot T_{rest} \tag{S6}$$

$$\frac{dT_{act}}{dt} = k_{act} \cdot f_{act} \cdot T_{rest} + k_{prolif} \cdot T_{act} \cdot \left(1 - \frac{T_{act} + T_{cyt}}{T_{act,max}}\right) - k_{diff} \cdot T_{act} - k_{exhaust} \cdot T_{act} \tag{S7}$$

$$\frac{dT_{cyt}}{dt} = k_{diff} \cdot T_{act} - k_{exhaust} \cdot T_{cyt} \tag{S8}$$

$$\frac{dT_{exh}}{dt} = k_{exhaust} \cdot (T_{act} + T_{cyt}) \tag{S9}$$

#### S1.5 Cytokine network

$$\frac{d[IFN\gamma]}{dt} = k_{prod,IFN\gamma} \cdot T_{cyt} \cdot f_{IL10} - k_{deg,IFN\gamma} \cdot [IFN\gamma] \tag{S10}$$

$$\frac{d[TNF\alpha]}{dt} = k_{prod,TNF\alpha} \cdot T_{cyt} \cdot f_{IL10} - k_{deg,TNF\alpha} \cdot [TNF\alpha] \tag{S11}$$

$$\frac{d[IL6]}{dt} = k_{prod,IL6,mac} \cdot f_{mac} \cdot mac_{pool,eff} \cdot f_{IL10} + J_{feedback} - k_{deg,IL6} \cdot [IL6] \tag{S12}$$

$$\frac{d[IL1\beta]}{dt} = k_{prod,IL1\beta,mac} \cdot f_{mac} \cdot mac_{pool,eff} \cdot f_{IL10} - k_{deg,IL1\beta} \cdot [IL1\beta] \tag{S13}$$

$$\frac{d[IL10]}{dt} = k_{prod,IL10,mac} \cdot f_{mac} \cdot mac_{pool,eff} - k_{deg,IL10} \cdot [IL10] \tag{S14}$$

where the STAT3 positive feedback term is:

$$J_{feedback} = k_{fb} \cdot \max([IL6] - [IL6]_{baseline}, 0) \cdot \frac{[IL6]}{K_{m,fb} + [IL6]} \cdot f_{mac} \quad \text{(S15)}$$

##### S1.6 Tocilizumab rescue modifications

At $t = t_{intervention}$ the system is modified: $J_{feedback} \to 0$ (complete IL-6R blockade abolishes STAT3 feedback) and $k_{deg,IL6} \to 0.3 \cdot k_{deg,IL6}$ (reduced receptor-mediated IL-6 clearance). All other equations remain unchanged.

##### S1.7 Initial conditions and numerical integration

Initial conditions: $T_{rest}(0) = 1{,}000$ cells/$\mu$L; $T_{act}(0) = T_{cyt}(0) = T_{exh}(0) = 0$; all cytokines at 0 pg/mL except $[IL6](0) = [IL6]_{baseline} = 5$ pg/mL; $C_{drug}(0) = $ dose$/V_{central}$ for bolus, 0 for infusion. ODEs were integrated with the LSODA adaptive-step solver (relative tolerance $10^{-8}$, absolute tolerance $10^{-10}$).

##### S1.8 Scope and modeling assumptions

Several structural properties should be noted. First, the resting T cell pool $T_{rest}$ only loses cells through activation and does not recover via homeostatic influx; dynamics are therefore valid for the acute CRS window (0 to 72 h) but do not capture recovery of the resting pool after CRS resolves. Second, because $T_{act}(0) = 0$, the logistic proliferation term is zero at $t = 0$ and becomes significant only after initial activation flux. Third, $f_{mac}$ is treated as a quasi-static function of instantaneous cytokine concentrations rather than a dynamic state; this captures steady-state amplification but not the slower kinetics of monocyte recruitment from bone marrow. Fourth, the cytokine network operates on plasma concentrations only; no tissue-specific compartments are modeled.

#### S2 Exploratory CAR-T Transfer Test

This section reports an exploratory extension of the locked downstream cytokine network to tisagenlecleucel (CTL019/Kymriah) CAR-T cell therapy. It is reported in the Supplementary rather than the main paper because it is **not a blind validation**: CAR-T-specific upstream parameters were formulated with knowledge of the Teachey 2016 [10] cytokine data and adjusted iteratively until simulated cytokine magnitudes were broadly consistent with the reported clinical ranges. It is therefore a **post hoc exploratory transfer test** of the locked downstream cytokine network onto a new upstream paradigm, intended to identify failure modes and class-specific biology that the current architecture does not yet capture, not to add a fourth drug class to the main cross-drug claim.

##### S2.1 Motivation and scope

CAR-T differs from antibody-based bispecifics in fundamental ways: (i) there is no classical drug pharmacokinetics, as CAR-T cells themselves are the active species; (ii) in vivo expansion proceeds over days rather than hours, with a doubling time of approximately 0.78 days [11]; (iii) cytokine production per CAR-T cell is lower than for polyclonal T cell engagers because activation requires encounter with $CD19^+$ target cells rather than direct receptor cross-linking; and (iv) the response is naturally self-limiting as tumor antigen is depleted by CAR-T-mediated killing. The 17 cytokine network parameters remained *locked* at the blinatumomab-calibrated values (verified by the SHA-256 hash referenced in the main paper); only the upstream block was replaced.

##### S2.2 Cohorts

Two cohorts were considered, both receiving the CTL019 construct (anti-CD19, 4-1BB co-stimulation, lentiviral):

- **Pediatric ALL** (CHOP NCT01626495, $N = 39$, 11 with severe CRS).
- **Adult ALL/CLL** (PENN NCT02030847 and NCT01029366, $N = 12$, 3 with severe CRS).

Quantitative cytokine peak data were taken from Teachey et al. [10] Supplemental Tables 9 and 11. The two cohorts differed in patient age, lymphodepletion regimen, and disease burden, and showed substantially different clinical cytokine peak magnitudes.

##### S2.3 Modeling approach

The drug pharmacokinetic compartment was bypassed by setting clearance to zero, and the activated T cell compartment $T_{act}$ was seeded directly at $t = 0$ with the post-infusion CAR-T cell count (5 cells/$\mu$L pediatric, 15 cells/$\mu$L adult). The proliferation rate was set to $k_{prolif} = 0.042\ h^{-1}$ to match the Stein 2019 [11] doubling time. Carrying capacity $T_{act,max}$ was set to 1,500 cells/$\mu$L for pediatric and 5,000 cells/$\mu$L for adult. The differentiation rate $k_{diff}$ was set substantially lower than for bispecifics (0.002 $h^{-1}$ pediatric, 0.005 $h^{-1}$ adult) to reflect the antigen-specific, target-encounter-dependent nature of CAR-T cytokine release. All 17 cytokine network parameters remained unchanged from the blinatumomab-calibrated values.

##### S2.4 Results

Predicted peak cytokine concentrations for both cohorts are reported in Table S1 and Figure S1.

The exploratory transfer test produced two main observations. First, the locked downstream cytokine network reproduced IL-6 and IFN-$\gamma$ within the reported clinical ranges for both pediatric and adult cohorts (4/4 cases). The pediatric prediction (peak IL-6 = 205 pg/mL) lies within the clinical range of 12-1,137 pg/mL, and the adult prediction (peak IL-6 = 1,130 pg/mL) is close to the clinical median of 1,088 pg/mL and well within the clinical range of 5-8,399 pg/mL. This suggests that the IL-6/IFN-$\gamma$ portion of the downstream cytokine architecture has at least partial transferability beyond antibody-based T cell engagers.

Second, the test exposed two clear failure modes. (i) TNF-$\alpha$ was substantially over-predicted in both cohorts, by approximately 130-fold (pediatric) and 120-fold (adult). Clinical CAR-T CRS is characterized by surprisingly flat TNF-$\alpha$ responses (median 0.69-1.59 pg/mL severe),

**Table S1:** Exploratory CAR-T transfer test: predicted vs. observed peak cytokine concentrations for pediatric and adult cohorts (Teachey 2016 severe CRS, day 1-3 peak). CAR-T-specific upstream parameters were chosen with knowledge of the cytokine target ranges and this is not a blind validation.

| Cohort | Cytokine | Predicted (pg/mL) | Observed median (pg/mL) | Range (pg/mL) | Within range? |
|---|---|---|---|---|---|
| | IL-6 | 205 | 50.4 | 12.0-1,137 | **Yes** |
| Pediatric ($N = 39$) | IFN-$\gamma$ | 42 | 50.4 | 16.8-544 | **Yes** |
| | TNF-$\alpha$ | 93 | 0.69 | 0.40-2.01 | *No (over)* |
| | IL-6 | 1,130 | 1,088 | 5.29-8,399 | **Yes** |
| Adult ($N = 12$) | IFN-$\gamma$ | 262 | 1,449 | 1.61-1,621 | **Yes** |
| | TNF-$\alpha$ | 192 | 1.59 | 0.67-8.29 | *No (over)* |

Source for clinical values: Teachey 2016 [10] Supplemental Tables 9 and 11. 4 of 6 cytokine peaks fell within the reported clinical ranges (IL-6 and IFN-$\gamma$ for both cohorts). TNF-$\alpha$ was substantially over-predicted in both cohorts, identifying a class-specific failure mode of the locked network for antigen-specific cell therapy contexts.

in contrast to elevated TNF-$\alpha$ seen in TGN1412 and OKT3. The locked cytokine network, calibrated on bispecific T cell engagers, includes a strong macrophage-derived TNF-$\alpha$ component that does not match the CAR-T phenotype. (ii) The age effect was qualitatively captured but quantitatively under-predicted: predicted adult-to-pediatric ratios were 5.5× for IL-6 and 6.2× for IFN-$\gamma$, compared to clinical ratios of 21.6× and 28.7× respectively. The direction is correct but the magnitude is too small.

##### S2.5 Interpretation and implications

These two failure modes both point to the same missing biology: **antigen-dependent engagement**. CAR-T cytokine release depends on encounter with $CD19^+$ target cells, and the response is naturally self-limited as tumor antigen is depleted. The current model has no tumor burden state variable and no engagement-limited cytokine production, so it cannot capture (a) the natural off-switch that keeps clinical TNF-$\alpha$ flat, or (b) the burden-dependent severity that distinguishes adult from pediatric cohorts. In addition, the model predicts CAR-T cytokine peak around day 14 of the simulation window, whereas clinical CAR-T CRS typically peaks at day 5-10 post-infusion; this delayed time course is consistent with the absence of antigen-depletion dynamics. Adding a tumor antigen depletion mechanism is the most natural next step for a proper CAR-T model and is identified in the main paper Future directions as a priority.

The exploratory transfer test should be interpreted as evidence that the IL-6/IFN-$\gamma$ portion of the downstream architecture transfers partially to cell therapy, while TNF-$\alpha$ biology and time course are class-dependent and require modality-specific upstream extensions that are beyond the scope of the present paper.

#### S3 Local Accepted Parameter Ensemble (ABC) - Distributional Plots

This section provides the full distributional plots for the local accepted parameter ensemble described in the main paper Section "Local robustness across an accepted parameter ensemble." The ensemble was generated by uniform ±30% prior sampling around each calibrated cytokine

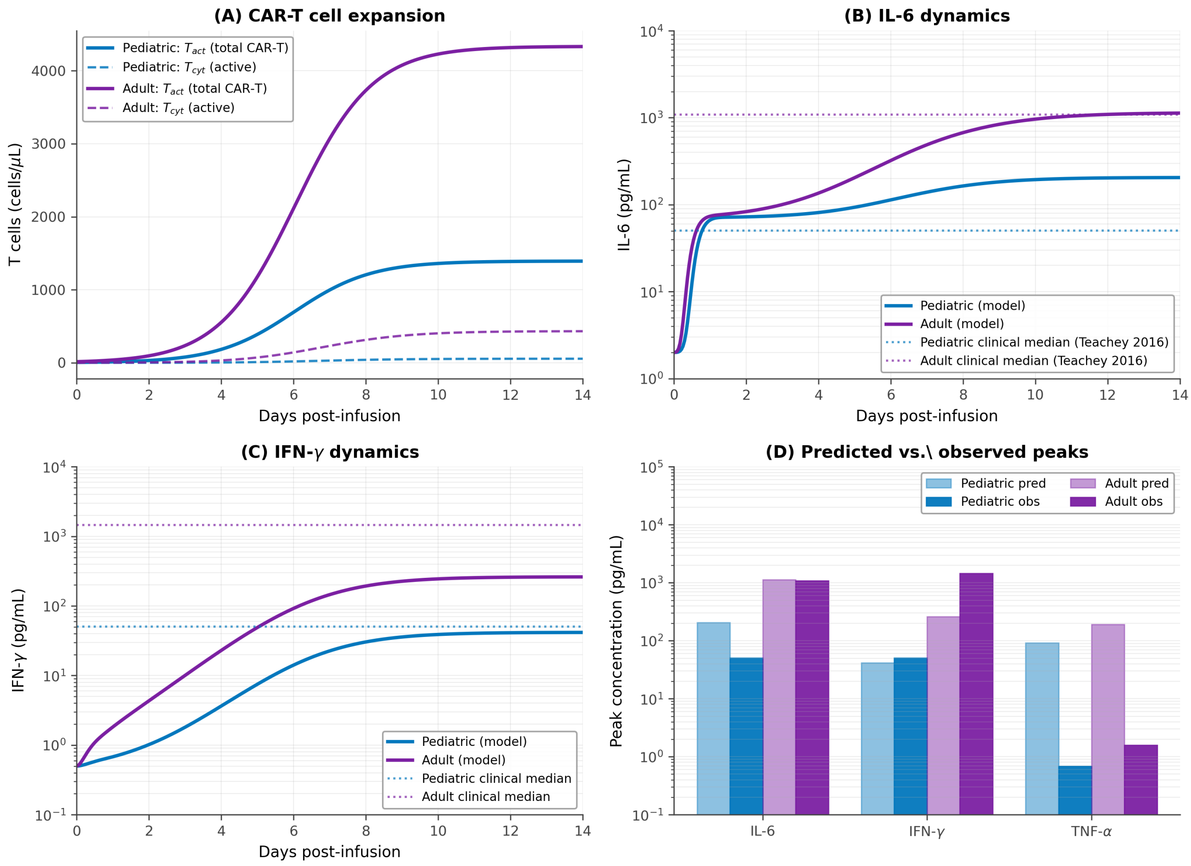


**Figure S1: Exploratory CAR-T transfer test (not a blind validation).** The locked downstream cytokine network was applied to a new CAR-T upstream module whose parameters were chosen with knowledge of the Teachey 2016 cytokine ranges. (A) Simulated CAR-T cell expansion: total CAR-T pool ($T_{act}$, solid) and active cytokine producers ($T_{cyt}$, dashed) for pediatric (dark blue) and adult (purple) cohorts over 14 days post-infusion, with doubling time matched to Stein 2019. (B) Simulated IL-6 dynamics with clinical median reference lines. Adult IL-6 peak (1,130 pg/mL) is very close to the clinical median (1,088 pg/mL, dotted); pediatric IL-6 peak (205 pg/mL) is within the reported clinical range of 12-1,137 pg/mL. (C) Simulated IFN-$\gamma$ dynamics, both within clinical ranges. (D) Predicted vs. observed peak cytokine concentrations (log scale). The downstream architecture transfers partially: IL-6 and IFN-$\gamma$ fall within clinical ranges for both cohorts, while TNF-$\alpha$ is consistently over-predicted, identifying a class-specific failure of the bispecific-calibrated network when applied to antigen-specific cell therapies.

network parameter, with an IL-6-only acceptance rule applied jointly across the three antibody-based drugs (blinatumomab, TGN1412, OKT3). Of 5,000 sampled parameter sets, 692 (13.8%) satisfied the joint acceptance criterion.

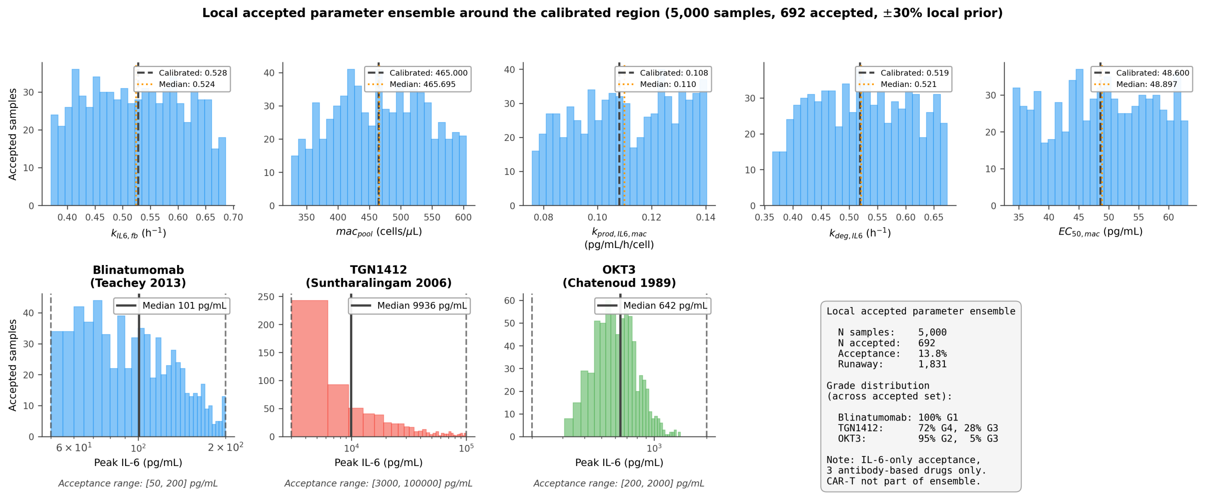


**Figure S2: Local accepted parameter ensemble around the calibrated region.** A local ABC-style ensemble (uniform ±30% prior around each calibrated value, IL-6-only acceptance rule, three antibody-based drugs only). (Top row) Distributions of the 5 free cytokine network parameters across the 692 accepted candidates from 5,000 samples. Black dashed line: calibrated value. Red dotted line: median across accepted samples. (Bottom row) Distributions of predicted peak IL-6 across the accepted ensemble for the three antibody-based drugs, with the a priori clinical acceptance ranges shown as black dashed lines. The accepted ensemble produces blinatumomab IL-6 in [55, 176] pg/mL (5-95%), TGN1412 IL-6 in [3,220, 71,677] pg/mL, and OKT3 IL-6 in [400, 1,012] pg/mL. Note: this is a local robustness ensemble characterizing the feasible region around the calibrated point on the IL-6 axis, not a full Bayesian posterior over the multi-cytokine network. CAR-T is not part of this analysis.

#### S4 Practical Identifiability of the Threshold-Setting Parameters

This section provides the constrained feasibility profile for the two threshold-setting parameters ($k_{IL6,fb}$ and $k_{deg,IL6}$) referenced in the main paper Section "Practical identifiability of the threshold-setting parameters."

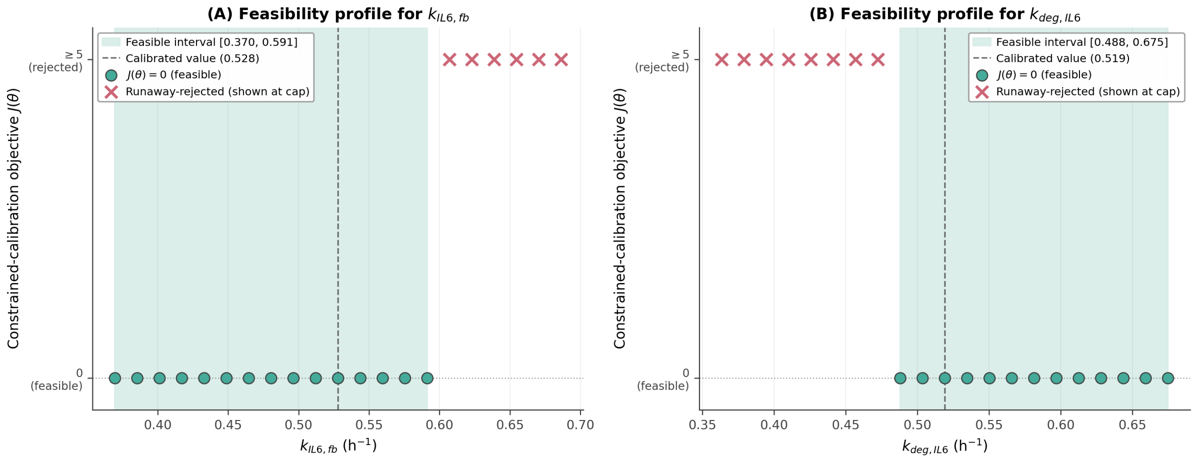


**Figure S3: Constrained feasibility profile for the two threshold-setting parameters.** This is *not* a classical smooth profile likelihood curve: because the calibration uses a range-based objective (zero within the clinical IL-6 ranges, $10^6$ hard-rejection if any drug triggers IL-6 runaway), each re-optimized grid point either returns $J(\theta) = 0$ (all three antibody-based drugs inside their clinical IL-6 ranges) or is rejected by the runaway criterion. Feasible points (teal circles) therefore sit on the $J = 0$ axis, while rejected points (rose-red crosses) are shown at a fixed visualisation cap of $J = 5$. Pale-teal band: contiguous feasible interval; dashed vertical line: calibrated value. (A) $k_{IL6,fb}$: the feasible interval spans $[0.370, 0.591]$ h$^{-1}$, with a sharp upper boundary where the feedback rate drives runaway amplification and a soft lower boundary because other parameters can compensate. (B) $k_{deg,IL6}$: the opposite asymmetry, with a sharp lower boundary at $\sim$0.488 h$^{-1}$ (below which IL-6 clearance is insufficient) and a soft upper boundary spanning to $\sim$0.675 h$^{-1}$. The complementary asymmetry indicates that neither parameter is individually practically identifiable; their combination encodes the IL-6 STAT3 amplification boundary and is identifiable as a *ratio* rather than individually.

### S5 Virtual Population Generator

The virtual population generator produced cohorts with physiologically correlated covariates. Demographics were sampled from sex-specific distributions: male height 176 ± 7 cm, female height 163 ± 6.5 cm, with weight correlated to height via allometric scaling and log-normal variability. BSA was computed using the Mosteller formula [8]. Estimated GFR was calculated using the CKD-EPI 2021 race-free equation [9]:

$$eGFR = 142 \cdot \min(SCr/\kappa, 1)^{\alpha} \cdot \max(SCr/\kappa, 1)^{-1.200} \cdot 0.9938^{age} \cdot S \qquad \text{(S16)}$$

with sex-specific $\kappa$ ($\kappa_F = 0.7$, $\kappa_M = 0.9$), $\alpha$ ($\alpha_F = -0.241$, $\alpha_M = -0.302$), and sex factor ($S_F = 1.012$, $S_M = 1.0$).

Four disease presets adjusted population demographics and biomarker distributions: healthy (default), generalized myasthenia gravis (gMG), B-cell ALL (B-ALL), and diffuse large B-cell lymphoma (DLBCL).

### S6 Full Sensitivity Analysis

One-at-a-time (OAT) parameter perturbation was performed for all four modules, varying each parameter by ±50% from its default value. The normalized sensitivity coefficient was $S = (\Delta Y/Y_0)/(\Delta p/p_0)$.

**Table S2:** OAT sensitivity coefficients (±50%) for key parameters across all modules.

| Module | Parameter | $S$ | Rank | Primary endpoint |
|---|---|---|---|---|
| IgG depletion | $FR_{IgG,baseline}$ | +0.56 | 1 | IgG nadir (%) |
| | $k_{deg,base}$ | −0.41 | 2 | |
| | $K_{d,IgG}$ | −0.25 | 3 | |
| | $K_{d,efg}$ | +0.20 | 4 | |
| Neutropenia | $slope$ | −3.72 | 1 | ANC nadir (cells/$\mu$L) |
| | $MTT$ | +1.45 | 2 | |
| | $ANC_{baseline}$ | +1.00 | 3 | |
| | $\gamma$ | +0.84 | 4 | |
| Hepatotoxicity | $k_{prime,decay}$ | −1.18 | 1 | Peak ALT (U/L) |
| | $P_{threshold}$ | −1.13 | 2 | |
| | $K_{m,conc}$ | −1.10 | 3 | |
| | $EC_{50,checkpoint}$ | −0.006 | 8 | |
| CRS | $mac_{pool}$ | +1.71 | 1 | Peak IL-6 (pg/mL) |
| | $k_{prod,IL6,mac}$ | +1.52 | 2 | |
| | $k_{IL6,fb}{}^{\dagger}$ | +2.91 | - | |
| | $EC_{50,mac}$ | −0.70 | 3 | |

$S = (\Delta Y/Y_0)/(\Delta p/p_0)$; $|S| > 1$ indicates output changes more than the input.
$^{\dagger}$ $k_{IL6,fb}$ sensitivity computed at ±20% due to bifurcation at larger perturbations.

### S7 Parameter Calibration Procedure

Parameters were calibrated using a sequential, module-by-module approach.

**IgG depletion.** Physiological parameters (compartment volumes, lymph flows, reflection coefficients) were taken from Shah and Betts [1]; drug-specific binding constants from Ulrichts and colleagues [2]; remaining parameters ($CL_p$, $CL_{renal}$, $FR_{IgG,baseline}$) were manually tuned to minimize AAFE against the single-dose Ulrichts 2018 dataset, then evaluated prospectively against multi-dose and ADAPT datasets.

**Neutropenia.** $MTT$, $\gamma$, and $slope$ were calibrated against Friberg 2002 docetaxel ANC time-course by manual iterative fitting. The recalibrated $\gamma = 0.214$ (vs. original 0.161) reflects use of effective one-compartment PK.

**Hepatotoxicity.** $P_{threshold}$ and $K_{m,conc}$ were calibrated to reproduce the 14% any-grade ALT elevation rate from CheckMate-067 [5].

**CRS.** The CRS module calibration proceeded in two stages. First, an initial parameter set was chosen by manual adjustment to bring simulated blinatumomab peak IL-6 into the Grade 1-2 range (50-200 pg/mL) reported by Teachey and colleagues [7]. The macrophage-gated STAT3 feedback structure was developed iteratively using TGN1412 as a model development case (see main manuscript Section 2.4). Second, the initial manual parameterization was refined using a **constrained blinatumomab calibration with soft cross-drug architectural preservation constraints** (main manuscript Section 2.7). The objective is range-based: it returns the squared log-fold distance to the nearest boundary if a drug prediction lies outside its published clinical IL-6 range, and zero otherwise. The published clinical ranges used (a priori, not modified after observing results) are blinatumomab 50-200 pg/mL (Teachey 2013, Grade 1-2), TGN1412 3,000-100,000 pg/mL (Suntharalingam 2006), and OKT3 200-2,000 pg/mL (Chatenoud 1989, Abramowicz 1989, Ferran 1990). **No specific clinical grade is required for any**

**drug**; only that each prediction fall within the corresponding published range. In particular, TGN1412 is not forced to the Grade 4 threshold; clinical TGN1412 is described as "Grade 3-4 in all six volunteers" (Suntharalingam 2006), so any prediction in the 3,000-100,000 pg/mL range is acceptable. The optimization optimizes only the five most influential cytokine network parameters identified by the prior one-at-a-time sensitivity analysis ($k_{IL6,fb}$, $mac_{pool}$, $k_{prod,IL6,mac}$, $k_{deg,IL6}$, $EC_{50,mac}$); the other 12 cytokine network parameters and all drug-specific T cell parameters are held fixed. Optimization uses derivative-free Nelder-Mead with ±50% bounds. Hard rejection ($J = 10^6$) is applied if any drug undergoes IL-6 amplification runaway (peak IL-6 $> 10^6$ pg/mL). Starting from the initial manual values, the optimizer converges to $J(\theta_{opt}) = 0$ (all three drug predictions inside their clinical ranges) with all parameter changes below ±10%. A ±10% local robustness perturbation analysis confirmed that two of the five free parameters ($k_{IL6,fb}$ and $k_{deg,IL6}$) sit near the IL-6 STAT3 amplification boundary while the other three are locally insensitive.

The robustness of the calibrated point was further characterized by a local ABC-style accepted parameter ensemble (5,000 samples within a uniform ±30% local prior, range-based IL-6-only acceptance against the same three antibody-based clinical ranges, 692 accepted, 13.8% acceptance rate) and a 2D $k_{IL6,fb}$ vs $k_{deg,IL6}$ stability map (900 grid points, 122 all-in-range, 0 wrong-order). See main manuscript Sections 2.7, 3.5, 3.6 for full details and the published code for reproducibility. CAR-T does not enter any of these analyses; it appears only as a post hoc exploratory transfer test in main manuscript Section 3.7.

The other three modules (IgG depletion, neutropenia, hepatotoxicity) retain manual parameter adjustment as documented above; formal calibration of these modules is deferred to future work.
